# Pil-Chp orchestrates a multi-level regulatory system to control type IV pilus dynamics in *Pseudomonas aeruginosa*

**DOI:** 10.64898/2026.09.03.749161

**Authors:** Farhana Haque, Zil K. Modi, Lincon Mazumder, Ahmed O. Yusuf, Ria Raam, Matthias D. Koch

## Abstract

The cyclical extension and retraction dynamics of type IV pili (T4P) mediate critical virulence traits in many bacterial species, yet how these dynamics are regulated remain poorly understood. Here, by analyzing individual pilus dynamics across mutants covering the entire T4P system of *Pseudomonas aeruginosa*, we identified active pili in several mutants previously classified as non-piliated. Correlating these dynamics with intracellular cAMP levels, transcriptional profiles, and T4P protein abundances reveals how the chemosensory Pil-Chp system controls T4P fiber length, count, extension rate, and polarity in surface-naïve cells by modulating cAMP levels. Specifically, fiber length is tuned by modulating the effective association and dissociation rates of the extension ATPase PilB; fiber count is regulated by controlling machine abundance; and extension rate is limited by the abundance of the major pilin (PilA). Furthermore, we demonstrate that these regulatory mechanisms of T4P length and count differentially impact T4P-dependent functions: phage infection efficiency increases continuously with fiber count or length, whereas twitching motility exhibits step-like activation thresholding. This indicates that the functional output of T4P dynamics is strictly context dependent. Because the Pil-Chp system is conserved across diverse human and plant pathogens, this multi-level regulatory logic may represent a widespread mechanism controlling T4P-mediated virulence.

## Introduction

Type IV pili (T4P) are virtually universal surface appendages found across both bacteria and archaea [1]. T4P mediate a diverse array of functions including adhesion, twitching motility, biofilm formation, DNA uptake, and virulence [2]. While the core assembly machinery is evolutionarily conserved, regulatory strategies are highly variable and often tailored to specific phylogenetic clades - such as the T4Pa, T4Pb, and Tad (T4Pc) systems - reflecting adaptations to diverse ecological niches [1, 3–7] .

Cyclic di-GMP (c-di-GMP) for example has been shown to modulate pilus activity in both Gram-negative and some Gram-positive species [8–10]. It acts directly by binding the extension ATPase MshE in *Vibrio cholerae* [8, 11], indirectly via c-di-GMP effector proteins (including PilZ-domain proteins, FimW, and FimX) to modulate the extension ATPase PilB in *Pseudomonas aeruginosa* [9, 12–16], and transcriptionally via a riboswitch that upregulates the major pilin gene *pilA1* in *Clostridium difficile* [17, 18]. Other common strategies include two-component systems such as PilSR, which senses inner-membrane pilin levels and regulates *pilA* transcription in many T4Pa pili-producing Proteobacteria [19–22], as well as orphan response regulators like DrdR, which directly binds the ATPases and regulates their activities in *Xanthomonas campestris* [23]. Dedicated chemosensory systems are also frequently employed to control motor activity and polarity, for example during motility and predation in *Myxococcus xanthus* [24–28] and *Pseudomonas aeruginosa* [29–32] or in cyanobacteria such as *Synechocystis sp*. PCC6803 during phototaxis [33, 34]. Interestingly, several bacteriophages exploit these T4P regulatory pathways as infection exclusion mechanisms. For instance, phages LUZ19 and JBD26 use PilZ to inhibit pilus extension, while phage PIT4 targets PilS to suppress major pilin production [35–37].

In contrast, the Pil-Chp chemosensory system constitutes a specialized surface-sensing module that is largely restricted to γ-Proteobacteria, but includes many important human (*Pseudomonas*, *Acinetobacter*) and plant (*Xanthomonas*, *Xylella*) pathogens [22, 38–41]. In *P. aeruginosa*, Pil-Chp integrates mechanosensing via the methyl-accepting chemotaxis protein PilJ with intracellular signaling through the hybrid kinase ChpA [15, 42–47]. ChpA autophosphorylates at multiple histidine phosphotransfer domains and transfers the phosphoryl group to the CheY-like response regulators PilG and PilH [30, 48, 49]. Current models propose that phosphorylated PilG promotes recruitment and activity of the extension ATPase PilB at the leading pole during cell migration, whereas phosphorylated PilH antagonizes PilG function (via phosphate-sink competition) to favor retraction and directional reversals [30, 42, 50]. In parallel, the phosphorelay stimulates the adenylate cyclase CyaB, elevating intracellular cAMP levels that activate the transcriptional regulator Vfr and its large regulon (>100 genes), many of which encode additional T4P components and virulence factors [43, 51, 52]. Accessory scaffold proteins such as FimL directly link components of the T4P alignment subcomplex PilMNOPQ (via FimV) to PilG, thereby coupling the pilus machinery to Pil-Chp signaling [41]. Crosstalk with c-di-GMP signaling pathways (e.g., involving PilY1 and the diguanylate cyclase SadC) further modulates the transition between acute and chronic infection lifestyles [12, 15, 45, 53].

However, while it is well established that Pil-Chp is important for T4P function, we still understand little about how Pil-Chp modulates the dynamics of individual T4P machines or how cAMP-dependent transcriptional changes are integrated with direct regulation of the motor ATPases by PilG and PilH. To address this gap, we directly quantified and correlated the dynamics of individual pili (extension frequency, fiber length, extension and retraction rates) in mutants spanning the entire T4P and Pil-Chp systems with corresponding changes in transcription and protein levels. These results are consistent with a model where Pil-Chp regulates the different levels of T4P dynamics through cAMP-dependent regulation of protein expression.

## Results

To better understand how Pil-Chp regulates pilus dynamics, we analyzed T4P activity in mutants of the entire T4P and Pil-Chp system (41 individual knock-out mutants) using Cysteine-maleimide labeling of pili in a *pilA*-A86C background [54–56].

### Direct fluorescent labeling reveals dynamic pilus extension in mutants previously thought to be non-piliated

First, we determined which strains assemble pilus fibers and observed visibly different degrees of piliation in the following mutants (see Figure 1A, Supplementary Movies 1-19): *pilH*, *pilK*, *chpB*, *cpdA* were hyper-piliated and made significantly more active pili than WT; *pilU*, *pilY2*, *chpE*, *fimL*, *fimT*, *fimX* had similar pilus activity to WT; and *pilG*, *pilI*, *pilJ*, *chpC*, *cyaB*, *fimV*, *vfr* had less pilus activity compared to WT. We note that all of these strains make dynamic pili except the deletion of the retraction motor *pilT* where pili are static. We further note that all mutants localized pili to the poles, except *fimV*, which is consistent with similar observations in *Acinetobacter baylyi* and *Pseudomonas* [57–59]. The following mutants did not make any visible pili in our experiments (see Supplementary Figure 1): *pilB*, *pilC*, *pilD*, *pilE*, *pilF*, *pilM*, *pilN*, *pilO*, *pilP*, *pilQ*, *pilR*, *pilS*, *pilV*, *pilW*, *pilX*, *pilY*1, *pilZ*, *chpA*, *chpD*, *fimS*, *fimU*, *fimW*. As most of these genes form structural components of either the T4P machine or the pilus fiber, these observations are consistent with previous reports and not surprising [60–63]. However, *chpA* and *fimW* have previously been reported to be piliated using Western blotting of sheared pili and electron microscopy, which our data of direct pilus labeling does not confirm [42, 64].

**Figure 1.**
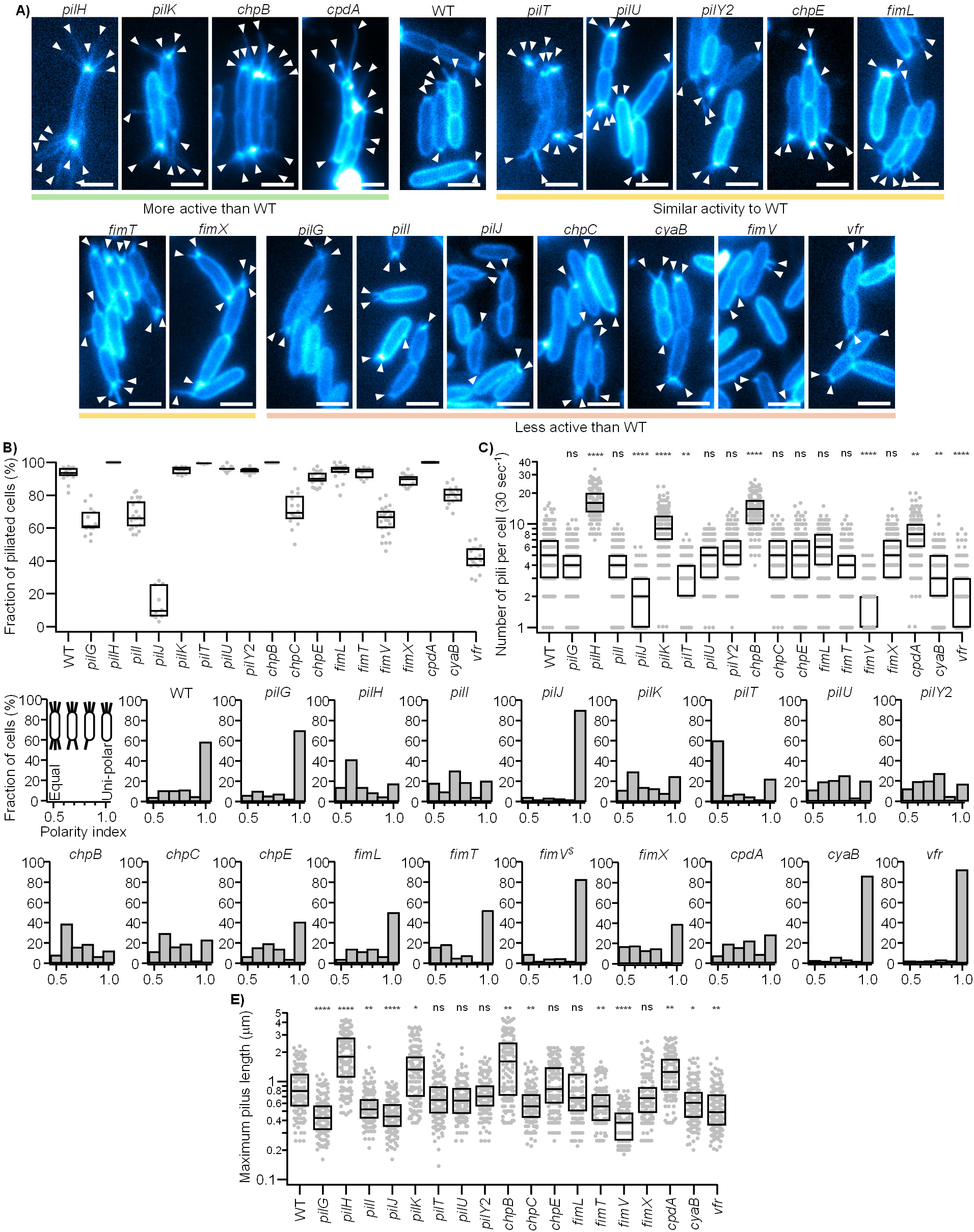
Impact of deletions in the T4P and Pil-Chp system genes on pilus dynamics in single cells. **A)** Representative still images of piliated mutants (see Supplementary Data for images of non-piliated mutants). Corresponding movies of dynamic pilus extensions and retractions are available as Supplementary Movies 1-19. White arrows point to individual pili. Scale bars: 2 μm. **B)** The fraction of cells generating visible pili. Data points represent individual movies of 30 second length. The fraction represents the ratio of individual cells that generated pili in each movie. N = 9 independent movies from 3 biological replicates (each with ≥3 technical replicates; 25–100 cells per movie). **C)** The number of pili individual cells generated in 30 second movies. N = 150 cells total from three biological replicates with 50 cells each were analyzed. **D)** Polarity of the pili extended by individual cells calculated as the count of pili on the dominant pole divided by the total number of pili for each cell. A polarity of 1 indicates all pili were made on just one pole, a polarity of 0.5 indicates both poles made an equal amount of pili. N = 150 cells from three biological replicates with 50 cells each were analyzed. $: While cells in the *fimV* mutant did not localize to the pole, individual pili were associated with the closest pole for the sake of analysis. **E)** The maximum length of individual pili. N = 150 pili from three biological replicates with 50 cells each were analyzed. **Statistical analysis:** Statistical significance indicates similarity to WT. B) Statistical significance was tested using a standard T-test of the mean of two distributions. C,E) Statistical significance was tested using a bootstrapped (10,000 iterations) non-parametric Wilcoxon-Mann-Whitney two-sample rank test: (ns) not significant, P > 0.05; (*) P < 0.05; (**) P < 0.01; (***) P < 0.001; (****) P < 0.0001. **Box plots:** Boxes represent median and interquartile range (25th–75th percentiles).

To quantify the degree of piliation in these mutants, we determined 1) the fraction of cells that made at least one pilus in 30 second long movies (out of approximately 50 cells per video) (see Figure 1B) and 2) the total number of pili that were made by piliated cells in the same 30 second time window (Figure 1C). While most mutants had a comparable degree of piliation to WT where ∼90% of cells were piliated and individual cells made an average five pili per 30 seconds, it is noteworthy that practically every single cell in the *pilH*, *cpdA*, and *chpB* backgrounds made pili with rare exceptions, and that the average cell in the these backgrounds (as well as *pilK*) made between 10 and 20 pili per 30 seconds, two- to four-times the WT extension frequency. Interestingly, while *pilG*, *pilI*, *pilJ*, *chpC*, *fimV*, *cyaB*, and *vfr* had a significantly smaller fraction of piliated cells compared to WT, many cells in these backgrounds still do extend one to two dynamic pili in the 30 second time windows analyzed (Figure 1C). This is surprising because most of these mutants (*pilG*, *pilI*, *pilJ*, *cyaB*) were previously believed to be non-piliated as they do not migrate in twitching assays and/or did not show bands in Western blots of sheared surface pili [42, 43]. This highlights that the presence of pili does not necessarily indicate biological function, and vice versa, that negative results in functional assays cannot be taken as strict proof for the absence of pilus fibers.

### Pil-Chp modulates the polarity of pilus extension

The analysis of T4P fiber dynamics also revealed clear mutant-dependent differences in the polarity of pilus extension, i.e., the number of pilus fibers on each pole of the cell (Figure 1D). We quantified this by calculating a polarity index, which is the number of pili on the dominant pole divided by the total number of pili in each cell. A ratio of one indicates all pili were made on one pole (zero on the other), while a ratio of 0.5 indicates equal amounts of pili on both poles (see first panel in Figure 1D). WT cells for example bias extension towards one pole with rare exemptions where cells make close to equal amounts of pili on both poles. Mutants that maintained this polarity were *pilG*, *pilJ*, *fimL*, *fimT*, *fimX*, *cyaB*, and *vfr*, while cells in *pilH*, *pilK*, *chpB*, and *cpdA* backgrounds generate close to equal amounts of pili on both poles. Interesting, pili in the *fimV* background localized to the side of the cell, but when associated with a pole by the minimal distance to either pole still maintained a polarity to one side of the cell. While these findings are largely consistent with recent models about how the Pil-Chp system orchestrates T4P polarity during surface interactions, our data indicate that these mechanisms regulate polarity even in the absence of a surface and reveal several previously unappreciated genes that affect pilus polarity, such as *pilI*, *pilU*, *pilY*2, *chpC*, and *chpE* [47, 50, 65, 66].

### Pil-Chp modulates the length of pilus fibers

After analyzing the general degree of piliation of individual cells, we next sought to determine how the length of pilus fibers is affected by mutations in the Pil-Chp system. While pilus length regulation is not well understood, the length of pilus fibers is crucial for the biological function of T4P [67]. If pili are not long enough to reach a surface for example, cells cannot migrate (twitching motility). For WT cells, forming the baseline, we found that the length of an average pilus fiber is ∼ 800 nm, consistent with prior data [56, 68, 69] (Figure 1E). Interestingly, we observed a similar pattern for pilus length between Pil-Chp mutants that we observed for the degree of piliation above: mutants like *pilG*, *pilI*, *pilJ*, *chpC*, *fimV*, *cyaB*, and *vfr* made significantly shorter (∼ 1.3-fold – 2-fold) pili, while *pilH*, *pilK*, *chpB*, and *cpdA* made significantly longer pili (1.5-fold – 2-fold). This demonstrates that Pil-Chp does not only regulate general activity of T4P, but also specific features of the dynamic cycle of pili.

### T4P activity is modulated by cAMP levels

Many of the mutants that displayed aberrant T4P activity have been implied in cAMP regulation and surface sensing, which is thought to form a transcriptional feedback with T4P biogenesis via Vfr [47, 51, 52, 70]. To better understand how this putative feedback mechanism affects pilus dynamics, we explicitly measured the cAMP levels in all mutant backgrounds that extended dynamic pili using the cAMP sensitive fluorescent PaQa reporter [47] (Figure 2A). We note that our data represents only the cAMP levels of liquid grown, surface-naïve cells and do not reflect surface sensing during prolonged surface exposure, in contrast to most studies [30, 45, 47]. This analysis reveals that the base level of cAMP in mutants of the T4P and Pil-Chp system varies by two orders of magnitude. Specifically, *pilH*, *pilK*, *chpB*, *fimT*, *fimX*, and *cpdA* had significantly elevated cAMP levels (2-20x), while *pilG*, *pilI*, *pilJ*, *pilY2*, *chpC*, *chpE*, *fimL*, *fimV*, *cyaB*, and *vfr* had significantly lower baseline levels of cAMP (2-5x) in surface-naïve cells compared to WT. These results are large consistent with previous β-galactosidase activity assays [43].

**Figure 2.**
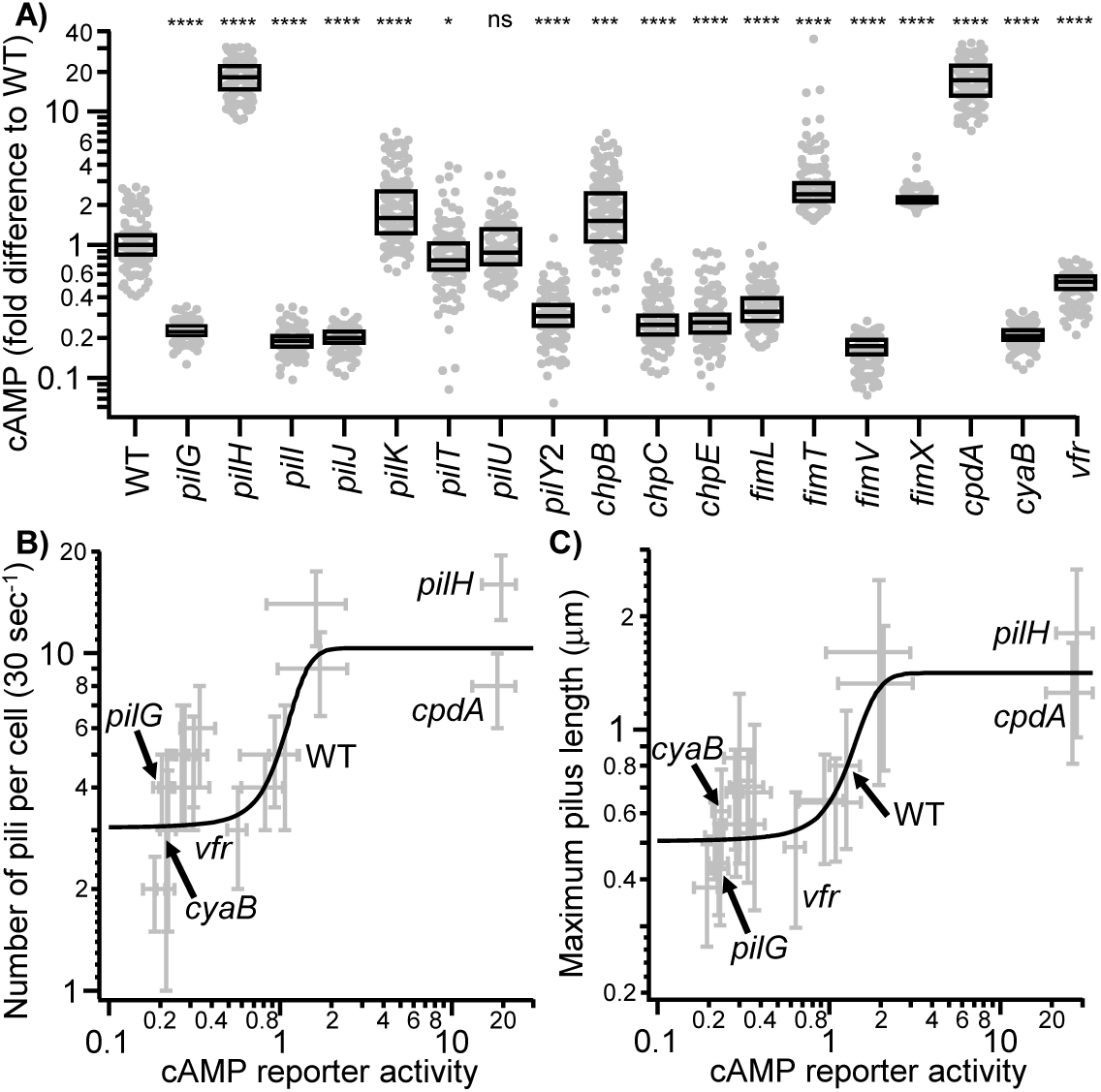
cAMP tunes the activity of T4P. **A)** cAMP levels in individual surface-naïve cells measured by the cAMP sensitive PaQa fluorescent transcriptional reporter. All data were normalized to the median cAMP level in WT. **B)** Correlation between the number of pili (from Figure 1C) and cAMP levels. **C)** Correlation between the length of individual pili (from Figure 1E) and cAMP levels. **Statistical analysis:** A) Statistical significance indicates similarity to WT and was tested using a bootstrapped (10,000 iterations) non-parametric Wilcoxon-Mann-Whitney two-sample rank test: (ns) not significant, P > 0.05; (*) P < 0.05; (**) P < 0.01; (***) P < 0.001; (****) P < 0.0001. B,C) Error bars indicate the inter quantile range around the medians. **Box plots:** Boxes represent median and interquartile range (25th–75th percentiles).

A linear correlation test between cAMP levels and the number of pili per cell (R = 0.65, P < 0.01) and the length of pili (R = 0.69, P < 0.001) reveals a positive correlation between cAMP levels and T4P activity. Even more so, correlation plots of both parameters as a function of the measured cAMP levels are described well by a Sigmoidal fit function (with reduced Chi^2^ = 1.37 for T4P number and 0.39 for pilus length), as expected for the cooperative activation of Vfr by cAMP (Figure 2B,C). The WT cAMP levels fall just below the middle of the steep transition region of the fitted sigmoidal curve where cAMP-dependent effects on T4P activity are most sensitive. Interestingly, the transition area ranges from ∼ 0.5-fold to ∼2-fold the WT levels of cAMP. This suggests that Pil-Chp can efficiently tune T4P dynamics by triggering small changes of cAMP levels in the cell. Further, similarity of cAMP levels between WT and *pilT* suggest that retraction deficiency has no direct effect on transcriptional regulation of T4P in surface-naïve cells.

### Pil-Chp modulates pilus length by tuning how long the ATPases interact with the T4P machine

Intrigued by the question of how Pil-Chp could modulate the dynamics of T4P fibers via cAMP, we sought to better understand the molecular mechanism behind how Pil-Chp modulate T4P fiber length. Since length is the product of speed (extension rate) and time (duration of extension), we analyze both parameters for representative low and high cAMP mutants (Figure 3). The extension and retraction rates of mutants with low cAMP levels (*pilG*, *cyaB*) were significantly (1.3x – 2x) lower than WT (∼230 nm/sec extension, ∼390 nm/sec retraction), while high cAMP mutants (*pilH*, *cpdA*) remained similar or slightly elevated to WT. In contrast, the extension and retraction durations of individual pili in high cAMP mutants were significantly elevated (2.6x – 3x) while low cAMP mutants remained indistinguishable from WT.

**Figure 3.**
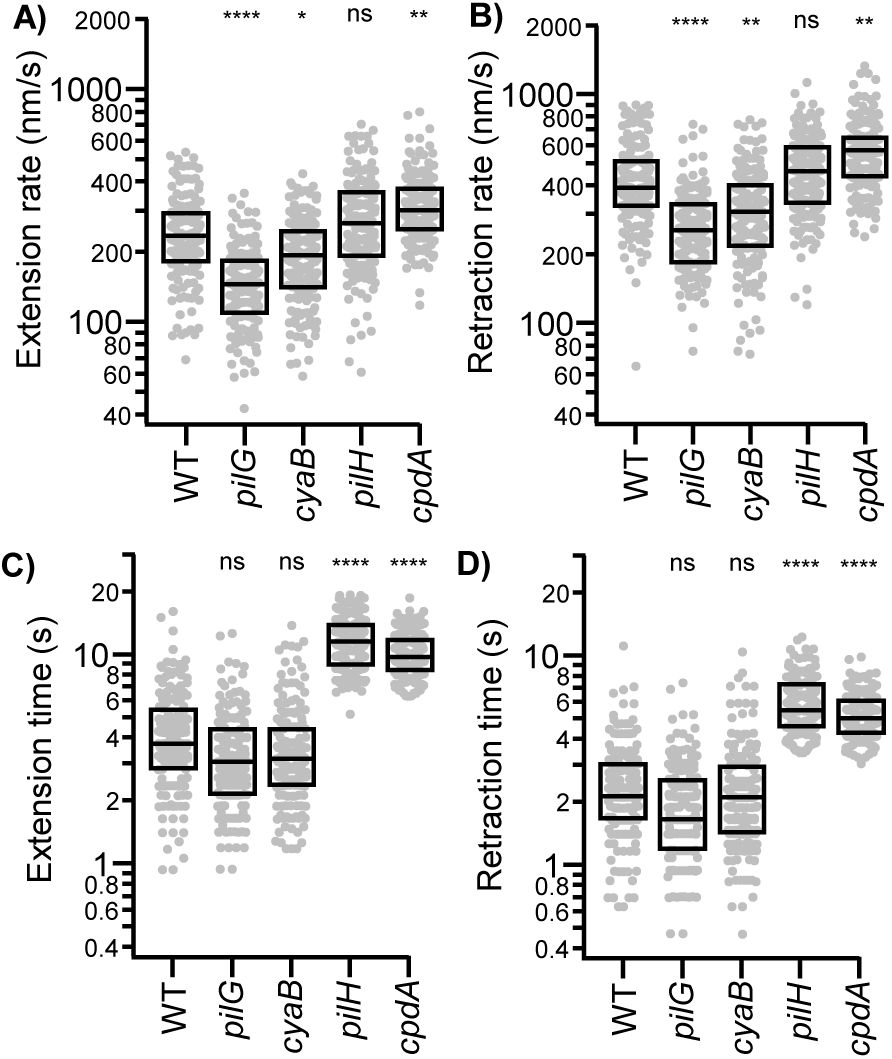
Pil-Chp modulates the extension/retraction rates and the total duration of extension/retraction of individual pilus fibers in mutants with low cAMP (*pilG*, *cyaB*) and high cAMP (*pilH*, *cpdA*). **A-D)** N = 150 cells from three biological replicates with 50 cells each were analyzed. **Statistical analysis:** Statistical significance indicates similarity to WT and was tested using a bootstrapped (10,000 iterations) non-parametric Wilcoxon-Mann-Whitney two-sample rank test: (ns) not significant, P > 0.05; (*) P < 0.05; (**) P < 0.01; (***) P < 0.001; (****) P < 0.0001. **Box plots:** Boxes represent median and interquartile range (25th–75th percentiles).

While these findings support proposed models where PilG and PilH modulate pilus dynamics by directly interacting with and regulating the extension and retraction ATPases [29, 42, 50], the similarity between *pilG* and the low cAMP adenyl-cyclase mutant *cyaB* as well as between *pilH* and the high cAMP phosphodiesterase mutant *cpdA* suggest that Vfr-dependent transcriptional feedback is a dominant factor in modulating pilus dynamics by Pil-Chp.

### Pil-Chp tunes T4P protein abundance via cAMP-dependent transcriptional regulation

To test the hypothesis that the observed changes in T4P dynamics in Pil-Chp mutants are driven by cAMP-dependent transcriptional changes, we sought to directly measure the promoter activity of key T4P operons using RNA-seq and compared the transcriptional profiles of WT cells to *pilG* (low cAMP) and *pilH* (high cAMP) mutants (Figure 4A). In the low cAMP background, transcript levels for both *pilA* and *pilB* are downregulated 2-fold, while the Pil-Chp operon (*pilIJ chpABCD*), the T4P machine operon (*pilMNOPQ*), the minor pilin operon (*fimU pilVWXY1Y2E*), and the two retraction motors *pilT* and *pilU* remained indistinguishable from WT levels. In the high cAMP background on the other hand, we observe significant upregulation of 2- to 9-fold for all seven T4P operons.

**Figure 4.**
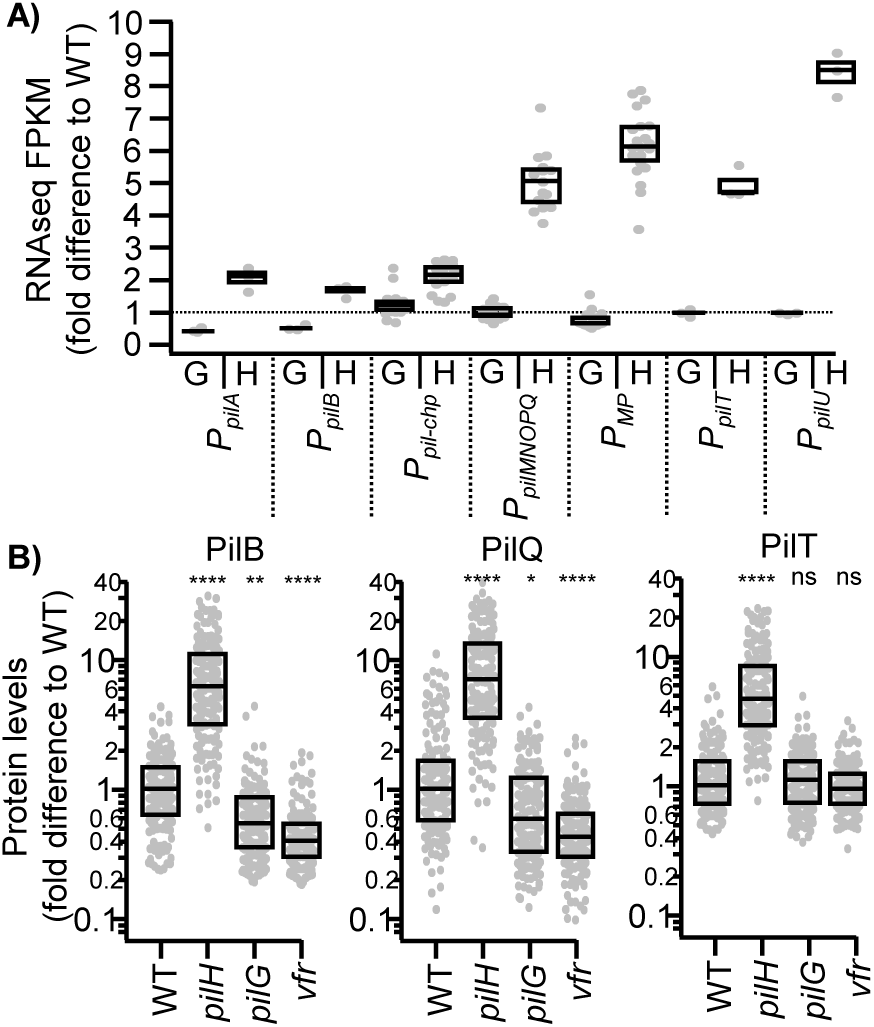
Transcriptional and translational regulation of T4P *genes* by Pil-Chp. **A)** Transcriptional levels of key T4P operons was measured using RNA-seq. Data points show the fold difference of *pilG* (G) and *pilH* (H) to WT. Data points for genes within the same operon were grouped as indicated (MP = minor pilins). N = 3 independent biological replicates per condition were analyzed. **B)** Protein levels of key T4P components were measured by transcriptional fusions and compared to WT. N = 150 cells from three biological replicates with 50 cells each were analyzed. **Statistical analysis:** Statistical significance indicates similarity to WT and was tested using a bootstrapped (10,000 iterations) non-parametric Wilcoxon-Mann-Whitney two-sample rank test: (ns) not significant, P > 0.05; (*) P < 0.05; (**) P < 0.01; (***) P < 0.001; (****) P < 0.0001. **Box plots:** Boxes represent median and interquartile range (25th–75th percentiles).

While this supports the hypothesis that Pil-Chp regulates T4P biogenesis, we wanted to confirm these results by measuring the levels of key T4P proteins to better understand how Pil-Chp regulates T4P dynamics. We chose to measure protein levels using fluorescent fusions to the motors PilB and PilT and the outer membrane pore PilQ as a representative T4P machine component. In accordance with the RNA-seq results, all three proteins showed a similar ∼5 to 7-fold increase in *pilH* vs WT. Further, PilB and PilQ showed slight but significant 2-fold decreases in *pilG* and *vfr*, while PilT did not. The results for PilA are largely consistent with prior reports that compared *pilA* transcription and PilA protein levels in *pilG* and *pilH* backgrounds [42]. Together, these data support a model where Pil-Chp regulates the expression of T4P genes to modulate T4P dynamics. For example, the generation of more T4P machines (via PilQ) in a high cAMP background explains the observation of more T4P extensions per time in *pilH* or *cpdA* mutants (see Discussion).

### The efficiency of phage infection gradually increases with increasing T4P count and fiber length

In Figure 1 we analyzed the detailed dynamics of pili of mutants of the entire T4P system. Since these mutants showed a large heterogeneity in T4P activity and dynamics, we sought to test how these changes affect the biological functions of T4P. T4P are common receptors for bacteriophages and little is known about how phage infection depends on changes in the number of T4P or their length. To address this gap, we performed a high throughput 96-well plate infection assay by measuring cell growth after infection with the T4P tip-protein specific phage JBD68 [67, 71]. As expected, we found that all strains that make active T4P show signs of reduced growth due to phage infection (see Supplementary Figure 2). However, we also found a clear pattern between pilus activity and the degree of cell growth: mutants with only few and short pili (e.g. *pilJ*) were only slightly impaired in growth and barely distinguishable from a *pilA* mutant control (Figure 5A), consistent with a low degree of infection efficiency. Cell growth decreased with increasing T4P activity (e.g. *pilG*) and reached WT-like behavior with a characteristic bump around 3-4 hours (given by initial cell growth), an intermediate decrease in growth afterwards (due to cell lysis by phage), and a subsequent rise after ∼10 hours due to the formation of lysogen. This phenotype was exaggerated by hyperactive T4P strains (e.g. *pilH*) which showed the strongest growth defect, consistent with the highest efficiency of phage infection. These results demonstrate that the efficiency of phage infection depends on and gradually changes with the frequency of pilus extension and the length of T4P fibers.

**Figure 5.**
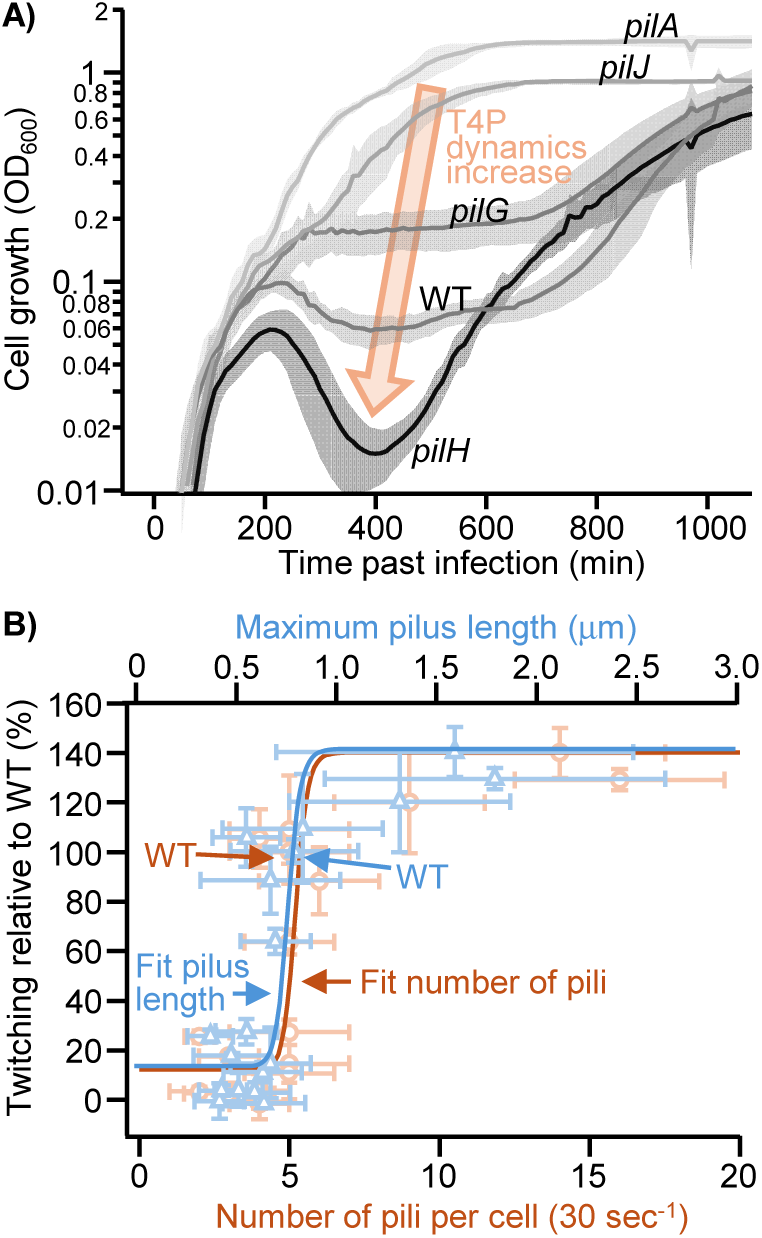
Changes in T4P activity tune biological functions of T4P such as phage infection and twitching motility. **A)** Growth curves of mutants with different T4P activity post infection (t = 0) with the T4P specific phage JBD68. **B)** Twitching motility activity as a function of the number of pili per cell and as a function of their length for all strains that made pili (excluding *cpdA* which was a sole outlier at 258% twitching).

### Surface-associated twitching motility undergoes a step-like functional transition with increasing T4P count and fiber length

Next, we sought to test how T4P activity impacts surface motility since this is commonly used to assay the activity of T4P and likely contributes to spreading in the host. To test this, we performed a stab agar twitch assay and compared twitch zones to WT and *pilA* controls after three days of incubation. As expected, mutants that did not make any visible pili (see Supplementary Figure 1) did not twitch (see Supplementary Figure 3) with the clear exception of *fimS* (and *fimW*) which was the second strongest twitching mutant in our assay (behind *cpdA*). How *fimS*, despite not making visible pili, could be such a strong twitching mutant is unclear, but likely a result of AlgR dependent exopolysaccharide regulation and/or upregulation of T4P activity during the three-day experiment due to mechanosensing [72]. Mutants that were classified as “less active than WT” in Figure 1A twitched significantly less (< 27%) than WT. Mutants that were classified as “similar activity to WT” display twitching between 64% to 109% compared to WT, with notable exception: *pilT* and *pilU* did not twitch, which is not surprising as these are the retraction motors, and *fimX* only twitched 15% of WT, which might be due to secondary effects of c-di-GMP regulation [16]. Lastly all mutants that were classified as “more active than WT” twitched above WT levels with 120% to 258% of WT.

Zooming out, our data show a clear correlation between T4P activity and twitching motility. To better understand this pattern, we plotted twitching motility as a function of the number of T4P and as a function of their length for mutants that made measurable pili (Figure 5B). Surprisingly, scatter plots of both relations show a high degree of similarity and overlap significantly. Correlation tests show R = 0.62 (P = 0.002, number of pili) and R = 0.73 (P = 0.0002, pilus length), supporting a strong correlation between twitching motility and T4P activity. Even more so, sigmoidal fits reveal a narrow transition region around WT T4P activity where twitching activity switches from zero to slightly above WT levels: mutants that make pili < 650nm average length do not twitch, while mutants making >850nm long pili twitch at a maximum rate of ∼140% of WT (∼800nm length) levels. Similarly, mutants making less than 5 pili do not twitch while mutants making more than 6 pili twitch at maximum rate. This narrow transition is surprising and suggests a high degree of optimization of the *Pseudomonas* T4P system for motility.

## Discussion

Here, we measured the detailed changes in pilus dynamics in individual cells across single-gene knockout mutants representing the entire extended type IV pilus (T4P) system in *Pseudomonas aeruginosa* (41 mutants). We believe this collection will serve as a valuable database for T4P dynamics for the community of T4P-related research. Our analysis revealed several mutants previously thought to be non-piliated that still extend dynamic pili. Furthermore, we quantified how baseline levels of cAMP in surface-naïve cells tune pilus dynamics. These findings support a multi-tiered molecular model for how Pil-Chp regulates T4P dynamics in *P. aeruginosa* (see Figure 6):

1. **Pil-Chp modulates T4P fiber count by regulating the expression of T4P machines:** The number of pilus fibers extended by a cell is constrained by the availability of functional T4P machines. We observed a strong dependence of pilus count on cAMP levels, characterized by reduced expression of T4P genes (such as the outer membrane pore protein PilQ) in low-cAMP backgrounds and increased expression in high-cAMP backgrounds. *pilH* cells exhibited ∼6-fold higher levels of PilQ, suggesting that cells generate ∼6-fold more T4P machines to support a higher frequency of pilus fiber extension. This increase in T4P machinery is further supported by a concurrent rise in the levels of both the extension and retraction ATPases (PilB and PilT, respectively) as well as the major pilin subunit PilA. Thus, our data suggests that Pil-Chp regulates the expression of T4P machinery to modulate total fiber count.
2. **Pil-Chp modulates pilus length primarily by regulating PilB expression:** Pilus length is determined by the extension rate and the duration of extension. Low-cAMP mutants displayed a slower extension rate but a similar extension duration, resulting in shorter pili [67]. Conversely, high-cAMP backgrounds exhibited ∼3-fold longer extension durations at similar extension rates, generating significantly longer pili. Higher expression of PilB in high-cAMP backgrounds aligns with this longer extension duration. Molecularly, PilB must assemble into an active hexameric complex to engage the T4P machinery and drive extension [73, 74] with dissociation or disassembly of the hexamer terminating the extension phase. Elevated cytosolic concentrations of PilB favor the assembly and structural stability of the functional hexameric ring, extending its operational lifetime at individual T4P machines before unbinding occurs to yield longer pilus fibers.
3. **Pil-Chp modulates the pilus extension rate by regulating PilA expression:** The extension rate driven by PilB is constrained by the intrinsic activity of the ATPase and the availability of the major pilin monomer PilA in the inner membrane [67]. In low-cAMP backgrounds, PilA expression decreased ∼2-fold. Lower levels of PilA force the extension ATPase into transient idle states: while the ATPase remains ready to process the next subunit, the reduced concentration of PilA requires more time for a subunit to diffuse to the T4P machinery [67]. This kinetic bottleneck reduces the overall extension rate. Importantly, cellular ATP concentrations in low-cAMP backgrounds did not differ significantly from wild-type (WT) levels, confirming that decreased extension rates stem from lower PilA concentrations rather than metabolic adaptation (see Supplementary Figure 4). Overexpressing *pilA* beyond WT levels (in high-cAMP backgrounds, e.g., *pilH*) did not significantly increase the extension rate and likely only supports the increased extension frequency of fibers per cell, suggesting that the effective ATP turnover rate of PilB is optimized for WT PilA concentrations.
4. **Pil-Chp modulates the polarity of T4P fibers:** Consistent with previous reports that Pil-Chp activates or deactivates T4P at the cell poles during surface sensing [30], we identified several mutants that disrupt typical WT polarity (predominantly unipolar) and yield mostly bipolar pili, even in surface-naïve cells. The precise molecular mechanisms governing polarity require further investigation. Notably, far more proteins are involved in maintaining polarity than previously recognized, including the auxiliary retraction ATPase PilU and several poorly characterized proteins such as PilY2, PilI, ChpC, and ChpE [66].

**Figure 6.**
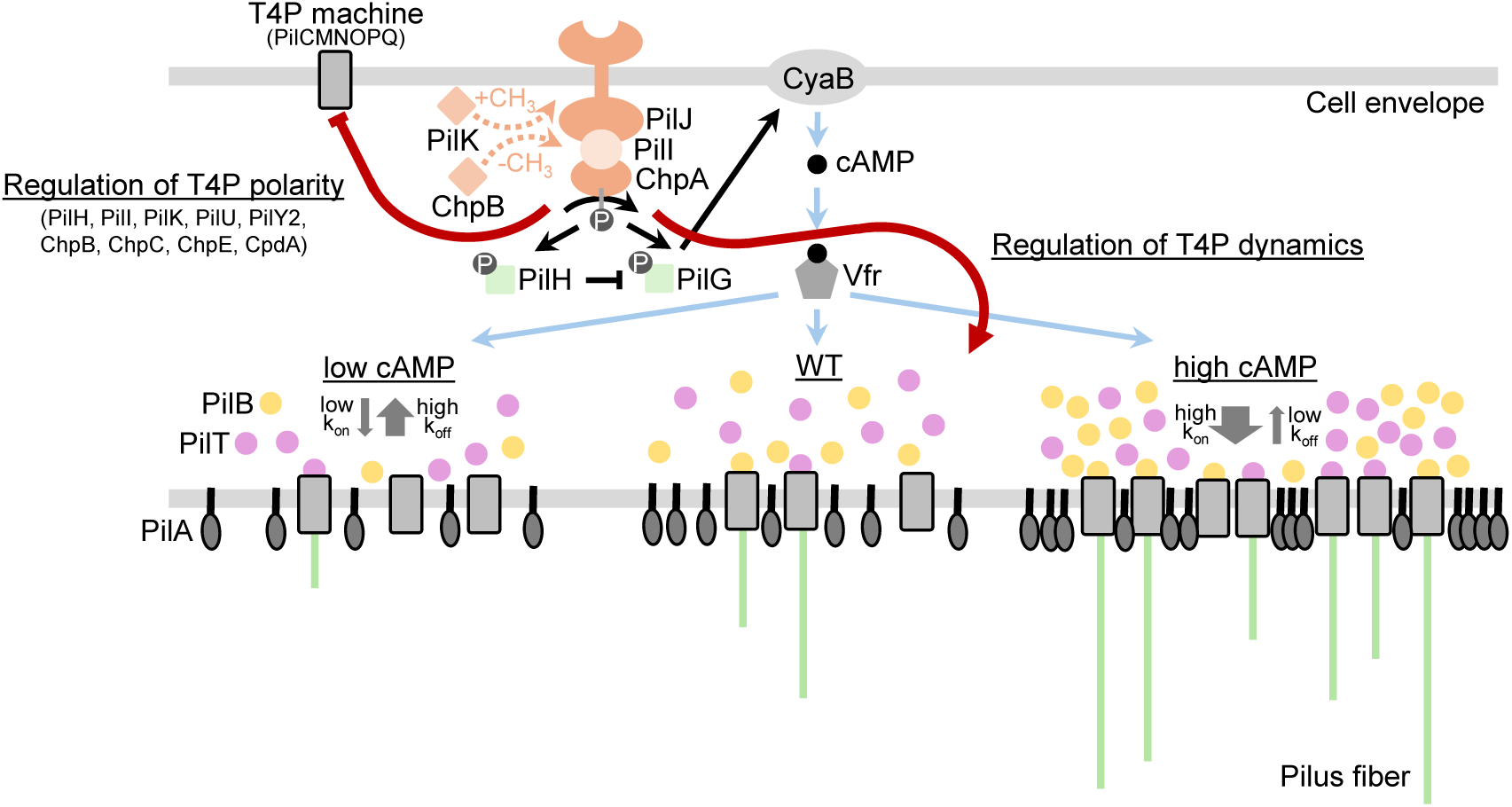
Schematic of the proposed molecular model of how Pil-Chp modulates T4P activity.

Together, these results suggest that *P. aeruginosa* leverages transcriptional feedback via Pil-Chp to regulate T4P dynamics across distinct, partially dependent processes. Our data from liquid-grown, surface-naïve cells establish the default ground state of T4P activity upon which surface contact engages the full PilG/PilH antagonism and dynamic PilK/ChpB localization to enable adaptive twitching [30, 50], effectively bridging adaptation between planktonic and surface-associated lifestyles.

While these findings demonstrate how Pil-Chp modulates T4P activity, a critical question remains: how do changes in T4P dynamics translate to alterations in T4P-dependent biological functions? We observed that phage infection efficiency scales gradually with T4P number or length, whereas twitching motility displays a sharp, threshold-like transition. This divergence illustrates that the relationship between T4P dynamics and functional outcomes varies depending on the specific biological process. For phages that bind the pilus tip, increasing pilus count or length elevates the probability of capturing a phage particle—explaining the gradual correlation between infection efficiency and T4P activity. For twitching motility, however, pili must be sufficiently long to reach the surface and pili below a minimal length threshold cannot contribute to motility [67]. Similarly, because the distribution of pilus number is exponential and skewed toward lower production rates, average pilus counts close to the minimum length required for surface interaction yields too few surface-interacting pili to support efficient motility. However, why the transition between non-motile and fully motile states is so narrow - and why additional or longer pili beyond this threshold do not enhance motility further - remains unclear.

Finally, a key question raised by our study is how biochemical changes in host environments modulate cAMP levels and basal T4P gene expression - both in liquid-like environments like the lung or bloodstream and during host surface contact in wounds or the urinary tract epithelium - to fine-tune T4P activity.

## Acknowledgements

We would like to thank Jessie Reed, Karen Maxwell, and Veronique Taylor for the gift of phage JBD68 and the entire Koch lab for stimulating discussion and the Department of Biology at Texas A&M for its supportive environment.

## Author contributions

F.H. and M.D.K. designed research and wrote the manuscript. F.H., Z.K.M., L.M., A.O.Y., R.R., and M.D.K. provided reagents, genetic constructs, or performed experiments and analyzed data.

## Data Availability

RNAseq data have been deposited in the NCBI Gene Expression Omnibus (GEO) under accession number GSE345958. Fluorescence microcopy movies of pilus dynamics are ∼500 GB total and available from the authors upon reasonable request. All other data needed to evaluate the conclusions in the paper are present in the paper and/or the Supplementary Materials.

## Methods and Protocols

### Construction of Strains

Knockout mutants and translational fusions in *Pseudomonas aeruginosa* PA01 were constructed using the two-step allelic exchange method [75]. For gene deletions, genomic sequences for target loci were retrieved from Pseudomonas.com and flanking upstream (P1/P2) and downstream (P3/P4) regions (∼500 bp) were amplified by PCR from chromosomal DNA. For the *pilQ* translational fusion, we amplified 500 bp flanking fragments using primers pilQ_FTL1_CT_F1.FOR/REV, pilQ_FTL1_CT_F2.FOR/REV from chromosomal DNA, and amplified the mScarlet-i3 fluorophore sequence from codon optimized cDNA (IDT) using primers FTL1_FP.FOR/REV, whereas *pilB* and *pilT* fluorescent fusion constructs were used from previous studies [56, 76]. For all constructs, the fragments were joined with the pEXG2 cloning vector backbone (digested with HindIII) using NEBuilder HiFi DNA Assembly Master Mix (New England Biolabs) according to the manufacturer’s protocol or by overlap extension PCR (SOE-PCR) and ligation. Constructs were then electroporated into donor *E. coli* S17 cells, followed by transformant validation via colony PCR and Sanger sequencing using pEXG2_Ver1/Ver2 primers. Biparental conjugation was conducted by harvesting mid-log phase donor cells (OD_600_ = 0.5) alongside heat-conditioned *P. aeruginosa* recipient cells (grown overnight, diluted 1:2 in fresh LB, and heat-shocked at 42°C for 3 hours). The puddle was scrapped off, resuspended in 150 μL PBS, then spread onto a Vogel-Bonner Minimal Medium (VBMM) plate containing 30 μg/ml gentamycin and incubated for 24 hours at 37 °C. For counter-selection, we struck single colonies on No-Salt Lysogeny Broth (NSLB) plates supplemented with 15% sucrose at 30 °C for 24 hours, and successful double-crossover mutants were confirmed via PCR amplification with locus-specific flanking primers and Sanger sequencing.

The ATP biosensor and the PaQa reporter plasmid were extracted from *Escherichia coli* S17 donor cells using a Qiagen miniprep kit [47, 76]. For the ATP biosensor, site-specific integration at the chromosomal phage insertion site required the helper plasmid pTSN2, which was isolated identically for co-transformation. Recipient *P. aeruginosa* cells were cultured to late-log phase, harvested, and rendered electrocompetent via three sequential washes in ice-cold 300 mM sucrose. Electrocompetent recipient cells were co-transformed with pTSN2 and either pAY9 or pAY11, or PaQa via electroporation, followed by recovery in 1 mL Lysogeny Broth (LB) at 37 °C while shaking for 1.5 hours. Cells were then plated on selective media and incubated overnight at 37°C. Successful transformants were verified by colony PCR and Sanger sequencing, then stored at -80 °C in 50% (v/v) glycerol.

### Fluorescence Microscopy imaging and analysis

#### Pilus Labelling and Imaging Experiments

For imaging, overnight grown samples in the *pilA*-A86C background were diluted 1: 300 in EZ rich medium and grown to mid log phase [56]. 180 μL samples were labelled with Alexa-488 dye and kept on ice for 10 minutes. Excess dye was washed away by pelleting and resuspending the cells twice with fresh EZ rich medium. About 2 µL of cells were spread on an agarose pad, allowed to air-dry for 2-3 minutes and then added to a 35mm glass bottom Mattek dish for imaging under a Nikon Ti2 Epifluorescence microscope with a ×100/1.45 NA Ph3 objective lens and a Hamamatsu FusionBT camera at 488nm. T4P dynamics videos were recorded at an exposure time of 200ms, a frame rate of 5 Hz, for 30 s each.

Data analysis was performed manually using ImageJ as previously described [56]. In short, the number of T4P was measured by counting the individual pili visible on single cells in the frames of the microscopy video. For pilus length, we identified the maximum length of each pilus in the video and measured the length from the cell body to the tip of the pilus using the line tool. The extension time was determined by counting the number of frames from the start of pilus extension (first frame with a visible pilus) to the maximum length was reached. Retraction time was measured similarly. For velocities, we measured the length change over at least three frames.

#### Translational Fusion Imaging

Single-colony overnight cultures were back-diluted 1:100 into 1 mL of Lysogeny Broth (LB) and incubated at 37 °C with shaking at 200 rpm. When the cells reached mid-log phase, 2 µL cell aliquots were spread on agarose pads and imaged using phase-contrast and fluorescence microscopy (561 nm channel). Quantitative image analysis was conducted using ImageJ/Fiji, wherein fluorescence images were thresholded to segment and isolate the fusion protein signal, followed by measurement and plotting of the total fluorescent intensity in each spot (cell pole) using the raw integrated density values.

#### PaQa Reporter Imaging

Bacterial strains were cultured overnight in Lysogeny Broth (LB) supplemented with 100 µg/mL carbenicillin at 37 °C with shaking at 250 rpm, back diluted 1:333 into fresh LB medium and grown to an OD_600_ of 0.5. Aliquots (300 µL) were harvested by centrifugation at 13,000 rpm for 2 minutes, the supernatant was removed, and cell pellets were resuspended in 300 µL of fresh LB medium. A 1 µL sample of each resuspended culture was spotted onto the center of a 1% agarose pad, which was then inverted onto a glass-bottom dish for imaging of the PaQa reporter. Single-cell image segmentation was performed using a custom-written, threshold-based algorithm driven by signal from a constitutive fluorescence reporter (PropD::mKate2) [45], enabling mean background estimation and intensity quantification across both spectral channels. Reporter activity was defined as the ratio of YFP to mKate2 fluorescence intensity.

#### ATP Biosensor Imaging

The overnight grown cells were back diluted 1:100 into 1 mL of Lysogeny Broth (LB) with 100 µg/mL carbenicillin and incubated at 37 °C with shaking at 200 rpm. When the cells reached mid-log phase, 2 µL cell aliquots were spread on agarose pads, covered with coverslips, and imaged using a Nikon Ti2 epifluorescence microscope equipped with a 100×/1.45 NA Ph3 objective lens. Dual-channel fluorescence imaging was conducted using 405 nm and 488 nm excitation wavelengths, respectively. Background-subtracted mean fluorescence intensities of individual cells were measured for each channel using the line scan function in ImageJ/Fiji, and ratio metric biosensor activity (405/488) was calculated and plotted as previously described [76, 77].

### Growth Assay of Phage Infection

Phage infection assays were performed in a 96-well liquid culture format as previously described [67]. Bacterial strains were cultured overnight in Lysogeny Broth (LB) at 37 °C with shaking at 250 rpm, back diluted 1:333 into 2 mL of fresh LB medium and grown at 37 °C with shaking to mid-log phase (OD_600_ = 0.5). Mid-log cultures were diluted 1:100 into fresh LB to achieve a working concentration of 5 × 10^6^ CFU/mL. For the infection assay, 25 µL of phage dilution (3.87 × 10^6^ PFU) was added to designated wells of a clear, flat-bottom 96-well microtiter plate, followed by 225 µL of diluted bacterial culture (1.12 × 10^6^ CFU total), bringing the final volume to 250 µL per well at a multiplicity of infection (MOI) of ∼3.44. Bacterial growth kinetics were continuously monitored by measuring OD_600_ every 10 minutes overnight at 37 °C with continuous orbital shaking in a microplate reader.

### Twitch Plate Assay

Twitching motility assays were performed using 1% (w/v) Lysogeny Broth (LB) agar plates, which were solidified and dried overnight at room temperature. Bacterial strains recovered from frozen glycerol stocks on standard 1.5% (w/v) LB agar plates at 37 °C were stab-inoculated using single colonies through the 1% agar layer to ensure contact with the bottom agar-plastic interface. Plates were incubated at 37 °C for 3 days to allow interstitial twitching motility zone expansion. Following incubation, the agar matrix was carefully removed, and the adherent biomass on the Petri dish surface was stained with 1% (w/v) crystal violet solution for 10–15 minutes, gently rinsed with deionized water, and air-dried. The surface area of each twitching motility zone (*T*_M_) was quantified by manually fitting a circle to images of the twitch zone, and relative motility was calculated by normalizing values against the wild-type strain (*T*_WT_, positive control) and a non-motile Δ*pilA* mutant (*T_pilA_* , negative control) using the formula 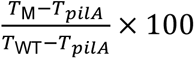 [67].

### RNA extraction and Sequencing

Total RNA was extracted by using the Promega Kit following the manufacturer protocol. To extract RNA, overnight culture of WT, *pilH* and *pilG* strains were diluted as 1:333 in fresh media and grown to OD_600_ of 0.8–1.0 before pelleting 1.0 mL of culture at 14,000 × g for 2 minutes and thoroughly removing the supernatant. The dry pellet was resuspended in 100 µL of fresh TE buffer containing lysozyme (0.4 mg/mL), tapped gently to mix, and incubated at room temperature for 3–5 minutes. Then 75 µL RNA Lysis Buffer was added followed by 350 µL RNA Dilution Buffer and mixed by inversion, strictly avoiding any centrifugation at this stage to prevent precipitating and losing the RNA. Next, 200 µL of 95% ethanol was mixed into the lysate by pipetting, transferred into a Spin Column Assembly, centrifuged at 12,000–14,000 × g for 1 minute. The flow-through was discarded and an initial wash with 600 µL ethanol-supplemented RNA Wash Solution (centrifuging again at 12,000–14,000 × g for 1 minute) was performed. For on-column DNase treatment, 40 µL Yellow Core Buffer, 5 µL 0.09M MnCl_2_, and 5 µL DNase I enzyme per sample were freshly mixed on ice, applied 50 µL directly onto the membrane, incubated at 20–25°C for 15 minutes, then 200 µL DNase Stop Solution was added, and centrifuged for 1 minute. The process was completed by washing with 600 µL RNA Wash Solution (1 minute spin), emptying the tube, performing a final dry wash spin with 250 µL RNA Wash Solution at high speed for 2 minutes, and transferring the spin basket into a fresh 1.5 mL Elution Tube. Rna was eluted by applying 100 µL Nuclease-free water across the membrane, centrifuging at 12,000–14,000 × g for 1 minute with the elution tube lids facing outward, discarding the basket, and storing the purified RNA at –70°C.

RNA was further processed for rRNA depletion using the Qiagen FastSelection kit and TruSeq library preparation and sequenced on an Illumina NovaSeq platform by the Texas A&M Institute for Genome Sciences and Society (TIGSS) center to generate 150 bp paired end reads. 8RNA-seq data were processed using the Galaxy cluster provided by the Texas A&M HPRC resources. Sequence alignment with the PAO1 reference genome was performed using HISAT2 and differential gene expression analysis was performed using DESeq2 to generate normalized FPKM values.

## References

1. Pelicic, V., Type IV pili: e pluribus unum? Molecular Microbiology, 2008. 68(4): p. 827–837.

2. Ellison, C.K., G.B. Whitfield, and Y.V. Brun, Type IV Pili: dynamic bacterial nanomachines. FEMS Microbiol Rev, 2022. 46(2).

3. Berry, J.L. and V. Pelicic, Exceptionally widespread nanomachines composed of type IV pilins: the prokaryotic Swiss Army knives. FEMS Microbiol Rev, 2015. 39(1): p. 134-54.

4. Denise, R., S.S. Abby, and E.P.C. Rocha, Diversification of the type IV filament superfamily into machines for adhesion, protein secretion, DNA uptake, and motility. PLOS Biology, 2019. 17(7): p. e3000390.

5. Giltner, C.L., Y. Nguyen, and L.L. Burrows, Type IV pilin proteins: versatile molecular modules. Microbiol Mol Biol Rev, 2012. 76(4): p. 740–72.

6. Makarova, K.S., E.V. Koonin, and S.-V. Albers, Diversity and Evolution of Type IV pili Systems in Archaea. Frontiers in Microbiology, 2016. **Volume** 7 **-** 2016.

7. Roberge, N.A. and L.L. Burrows, Building permits-control of type IV pilus assembly by PilB and its cofactors. J Bacteriol, 2024. 206(12): p. e0035924.

8. Floyd, K.A., et al., c-di-GMP modulates type IV MSHA pilus retraction and surface attachment in Vibrio cholerae. Nat Commun, 2020. 11(1): p. 1549.

9. Jain, R., et al., Type IV pilus assembly in Pseudomonas aeruginosa over a broad range of cyclic di-GMP concentrations. J Bacteriol, 2012. 194(16): p. 4285-94.

10. Zhang, X., et al., A comprehensive review of the pathogenic mechanisms of Pseudomonas aeruginosa: synergistic effects of virulence factors, quorum sensing, and biofilm formation. Front Microbiol, 2025. 16: p. 1619626.

11. Jones, C.J., et al., C-di-GMP Regulates Motile to Sessile Transition by Modulating MshA Pili Biogenesis and Near-Surface Motility Behavior in Vibrio cholerae. PLoS Pathog, 2015. 11(10): p. e1005068.

12. Almblad, H., et al., The Cyclic AMP-Vfr Signaling Pathway in Pseudomonas aeruginosa Is Inhibited by Cyclic Di-GMP. J Bacteriol, 2015. 197(13): p. 2190-200.

13. Jain, R., O. Sliusarenko, and B.I. Kazmierczak, Interaction of the cyclic-di-GMP binding protein FimX and the Type 4 pilus assembly ATPase promotes pilus assembly. PLOS Pathogens, 2017. 13(8): p. e1006594.

14. Laventie, B.J., et al., A Surface-Induced Asymmetric Program Promotes Tissue Colonization by Pseudomonas aeruginosa. Cell Host Microbe, 2019. 25(1): p. 140–152.e6.

15. Luo, Y., et al., A hierarchical cascade of second messengers regulates Pseudomonas aeruginosa surface behaviors. mBio, 2015. 6(1).

16. Roberge, N., et al., Twitching motility suppressors reveal a role for FimX in type IV pilus extension dynamics. PLoS Genet, 2025. 21(10): p. e1011802.

17. Bordeleau, E., et al., Cyclic di-GMP riboswitch-regulated type IV pili contribute to aggregation of Clostridium difficile. J Bacteriol, 2015. 197(5): p. 819-32.

18. McKee, R.W., et al., Type IV Pili Promote Clostridium difficile Adherence and Persistence in a Mouse Model of Infection. Infection and Immunity, 2018. 86(5): p. 10.1128/iai.00943-17.

19. Hobbs, M., et al., PilS and PilR, a two-component transcriptional regulatory system controlling expression of type 4 fimbriae in Pseudomonas aeruginosa. Mol Microbiol, 1993. 7(5): p. 669–82.

20. Kilmury, S.L.N. and L.L. Burrows, The Pseudomonas aeruginosa PilSR Two-Component System Regulates Both Twitching and Swimming Motilities. mBio, 2018. 9(4).

21. Rijal, A. and P.D. Curtis, Type IV pilin regulation: a transcriptional overview. Crit Rev Microbiol, 2026. 52(1): p. 36–63.

22. Vesel, N. and M. Blokesch, Pilus Production in Acinetobacter baumannii Is Growth Phase Dependent and Essential for Natural Transformation. J Bacteriol, 2021. 203(8).

23. Ren, P., et al., Tandem receiver domain response regulator DrdR modulates swarming by controlling PilB/PilT activity in Xanthomonas campestris. mLife, 2026. 5(4): p. 473–485.

24. Inclán, Y.F., H.C. Vlamakis, and D.R. Zusman, FrzZ, a dual CheY-like response regulator, functions as an output for the Frz chemosensory pathway of Myxococcus xanthus. Mol Microbiol, 2007. 65(1): p. 90–102.

25. Kaimer, C. and D.R. Zusman, Phosphorylation-dependent localization of the response regulator FrzZ signals cell reversals in Myxococcus xanthus. Mol Microbiol, 2013. 88(4): p. 740–53.

26. Keilberg, D., et al., A response regulator interfaces between the Frz chemosensory system and the MglA/MglB GTPase/GAP module to regulate polarity in Myxococcus xanthus. PLoS Genet, 2012. 8(9): p. e1002951.

27. Mercier, R., et al., The polar Ras-like GTPase MglA activates type IV pilus via SgmX to enable twitching motility in Myxococcus xanthus. Proc Natl Acad Sci U S A, 2020. 117(45): p. 28366–28373.

28. Sun, H., D.R. Zusman, and W. Shi, Type IV pilus of Myxococcus xanthus is a motility apparatus controlled by the frz chemosensory system. Curr Biol, 2000. 10(18): p. 1143–6.

29. Kühn, M.J., et al., Mechanotaxis directs Pseudomonas aeruginosa twitching motility. Proc Natl Acad Sci U S A, 2021. 118(30).

30. Patino, R., et al., Antagonistic response regulators spatially regulate receptor methylation in the Pseudomonas aeruginosa Pil-Chp surface sensing system. Cell Rep, 2025. 44(4): p. 115536.

31. Yarrington, K.D. and D.H. Limoli, The type IV pilus steering committee: how Pil-Chp controls directional motility. J Bacteriol, 2025. 207(11): p. e0039624.

32. Yarrington, K.D., T.N. Shendruk, and D.H. Limoli, The type IV pilus chemoreceptor PilJ controls chemotaxis of one bacterial species towards another. PLoS Biol, 2024. 22(2): p. e3002488.

33. Bhaya, D., A. Takahashi, and A.R. Grossman, Light regulation of type IV pilus-dependent motility by chemosensor-like elements in Synechocystis PCC6803. Proc Natl Acad Sci U S A, 2001. 98(13): p. 7540-5.

34. Harwood, T.V., et al., The cyanobacterial taxis protein HmpF regulates type IV pilus activity in response to light. Proceedings of the National Academy of Sciences, 2021. 118(12): p. e2023988118.

35. Hendrix, H., et al., PlzR regulates type IV pili assembly in Pseudomonas aeruginosa via PilZ binding. Nat Commun, 2024. 15(1): p. 8717.

36. Schroven, K., et al., The phage-encoded PIT4 protein affects multiple two-component systems of Pseudomonas aeruginosa. Microbiol Spectr, 2023. 11(6): p. e0237223.

37. Taylor, V.L., et al., Prophages block cell surface receptors to preserve their viral progeny. Nature, 2025. 644(8078): p. 1049-1057.

38. Cursino, L., et al., Identification of an operon, Pil-Chp, that controls twitching motility and virulence in Xylella fastidiosa. Mol Plant Microbe Interact, 2011. 24(10): p. 1198–206.

39. Ellison, C.K., et al., Subcellular localization of type IV pili regulates bacterial multicellular development. Nat Commun, 2022. 13(1): p. 6334.

40. Whitchurch, C.B., et al., Pseudomonas aeruginosa fimL regulates multiple virulence functions by intersecting with Vfr-modulated pathways. Mol Microbiol, 2005. 55(5): p. 1357–78.

41. Inclan, Y.F., et al., A scaffold protein connects type IV pili with the Chp chemosensory system to mediate activation of virulence signaling in Pseudomonas aeruginosa. Mol Microbiol, 2016. 101(4): p. 590–605.

42. Bertrand, J.J., J.T. West, and J.N. Engel, Genetic analysis of the regulation of type IV pilus function by the Chp chemosensory system of Pseudomonas aeruginosa. J Bacteriol, 2010. 192(4): p. 994-1010.

43. Fulcher, N.B., et al., The Pseudomonas aeruginosa Chp chemosensory system regulates intracellular cAMP levels by modulating adenylate cyclase activity. Mol Microbiol, 2010. 76(4): p. 889–904.

44. Geiger, C.J. and G.A. O’Toole, Evidence for the Type IV Pilus Retraction Motor PilT as a Component of the Surface Sensing System in Pseudomonas aeruginosa. J Bacteriol, 2023. 205(7): p. e0017923.

45. Koch, M.D., et al., Pseudomonas aeruginosa distinguishes surfaces by stiffness using retraction of type IV pili. Proceedings of the National Academy of Sciences, 2022. 119(20): p. e2119434119.

46. Leech, A.J. and J.S. Mattick, Effect of site-specific mutations in different phosphotransfer domains of the chemosensory protein ChpA on Pseudomonas aeruginosa motility. J Bacteriol, 2006. 188(24): p. 8479–86.

47. Persat, A., et al., Type IV pili mechanochemically regulate virulence factors in Pseudomonas aeruginosa. Proceedings of the National Academy of Sciences, 2015. 112(24): p. 7563–7568.

48. Darzins, A., Characterization of a Pseudomonas aeruginosa gene cluster involved in pilus biosynthesis and twitching motility: sequence similarity to the chemotaxis proteins of enterics and the gliding bacterium Myxococcus xanthus. Mol Microbiol, 1994. 11(1): p. 137–53.

49. Silversmith, R.E., et al., Phosphoryl Group Flow within the Pseudomonas aeruginosa Pil-Chp Chemosensory System: DIFFERENTIAL FUNCTION OF THE EIGHT PHOSPHOTRANSFERASE AND THREE RECEIVER DOMAINS. J Biol Chem, 2016. 291(34): p. 17677-91.

50. Kühn, M.J., et al., Two antagonistic response regulators control Pseudomonas aeruginosa polarization during mechanotaxis. Embo j, 2023. 42(7): p. e112165.

51. Wolfgang, M.C., et al., Coordinate regulation of bacterial virulence genes by a novel adenylate cyclase-dependent signaling pathway. Dev Cell, 2003. 4(2): p. 253–63.

52. Inclan, Y.F., M.J. Huseby, and J.N. Engel, FimL Regulates cAMP Synthesis in Pseudomonas aeruginosa. PLOS ONE, 2011. 6(1): p. e15867.

53. Webster, S.S., et al., Force-Induced Changes of PilY1 Drive Surface Sensing by Pseudomonas aeruginosa. mBio, 2021. 13(1): p. e0375421.

54. Ellison, C.K., et al., Real-time microscopy and physical perturbation of bacterial pili using maleimide-conjugated molecules. Nat Protoc, 2019. 14(6): p. 1803–1819.

55. Ellison, C.K., et al., Obstruction of pilus retraction stimulates bacterial surface sensing. Science, 2017. 358(6362): p. 535-538.

56. Koch, M.D., et al., Competitive binding of independent extension and retraction motors explains the quantitative dynamics of type IV pili. Proceedings of the National Academy of Sciences, 2021. 118(8): p. e2014926118.

57. Ellison, C.K., et al., Acinetobacter baylyi regulates type IV pilus synthesis by employing two extension motors and a motor protein inhibitor. Nature Communications, 2021. 12(1): p. 3744.

58. Ellison, C.K., et al., Subcellular localization of type IV pili regulates bacterial multicellular development. Nature Communications, 2022. 13(1): p. 6334.

59. Kazmierczak, B.I., M.B. Lebron, and T.S. Murray, Analysis of FimX, a phosphodiesterase that governs twitching motility in Pseudomonas aeruginosa. Molecular Microbiology, 2006. 60(4): p. 1026–1043.

60. Buensuceso, R.N.C., et al., Cyclic AMP-Independent Control of Twitching Motility in Pseudomonas aeruginosa. Journal of Bacteriology, 2017. 199(16): p. 10.1128/jb.00188-17.

61. Burrows, L.L., Pseudomonas aeruginosa twitching motility: type IV pili in action. Annu Rev Microbiol, 2012. 66: p. 493–520.

62. Chiang, P., M. Habash, and L.L. Burrows, Disparate subcellular localization patterns of Pseudomonas aeruginosa Type IV pilus ATPases involved in twitching motility. J Bacteriol, 2005. 187(3): p. 829–39.

63. Marko, V.A., et al., Pseudomonas aeruginosa type IV minor pilins and PilY1 regulate virulence by modulating FimS-AlgR activity. PLOS Pathogens, 2018. 14(5): p. e1007074.

64. Laventie, B.-J., et al., A Surface-Induced Asymmetric Program Promotes Tissue Colonization by Pseudomonas aeruginosa. Cell Host & Microbe, 2019. 25(1): p. 140–152.e6.

65. Kühn, M.J., et al., Mechanotaxis directs Pseudomonas aeruginosa twitching motility. Proceedings of the National Academy of Sciences, 2021. 118(30): p. e2101759118.

66. Kline, A., et al., Distinct roles of PilI and ChpC in Pseudomonas aeruginosa Chp chemosensory array architecture. Applied and Environmental Microbiology. 0(0): p. e01286–26.

67. Modi, Z.K., et al., Type IV pilus length determines virulence by regulating a putative subpopulation of non-contributing filaments in Pseudomonas aeruginosa. Communications Biology, 2026.

68. Talà, L., et al., Pseudomonas aeruginosa orchestrates twitching motility by sequential control of type IV pili movements. Nature Microbiology, 2019. 4(5): p. 774–780.

69. Skerker, J.M. and H.C. Berg, Direct observation of extension and retraction of type IV pili. Proc Natl Acad Sci U S A, 2001. 98(12): p. 6901-4.

70. Kanack, K.J., et al., Characterization of DNA-binding specificity and analysis of binding sites of the Pseudomonas aeruginosa global regulator, Vfr, a homologue of the Escherichia coli cAMP receptor protein. Microbiology, 2006. 152(12): p. 3485–3496.

71. Harvey, H., et al., Pseudomonas aeruginosa defends against phages through type IV pilus glycosylation. Nat Microbiol, 2018. 3(1): p. 47–52.

72. Marko, V.A., et al., Pseudomonas aeruginosa type IV minor pilins and PilY1 regulate virulence by modulating FimS-AlgR activity. PLoS Pathog, 2018. 14(5): p. e1007074.

73. Chlebek, J.L., et al., PilT and PilU are homohexameric ATPases that coordinate to retract type IVa pili. PLOS Genetics, 2019. 15(10): p. e1008448.

74. McCallum, M., et al., The molecular mechanism of the type IVa pilus motors. Nature Communications, 2017. 8(1): p. 15091.

75. Hmelo, L.R., et al., Precision-engineering the Pseudomonas aeruginosa genome with two-step allelic exchange. Nat Protoc, 2015. 10(11): p. 1820–41.

76. Yusuf, A.O., Z.K. Modi, and M.D. Koch, Rapid activation of dormant type IV pili enables a dispersal–infection tradeoff in environments with fluctuating nutrients. bioRxiv, 2026: p. 2026.04.23.720421.

77. Yaginuma, H., et al., Diversity in ATP concentrations in a single bacterial cell population revealed by quantitative single-cell imaging. Sci Rep, 2014. 4: p. 6522.

