## Supplementary Information for "Pil-Chp orchestrates a multi-level regulatory system to control type IV pilus dynamics in *Pseudomonas aeruginosa*"

### Supplementary Data

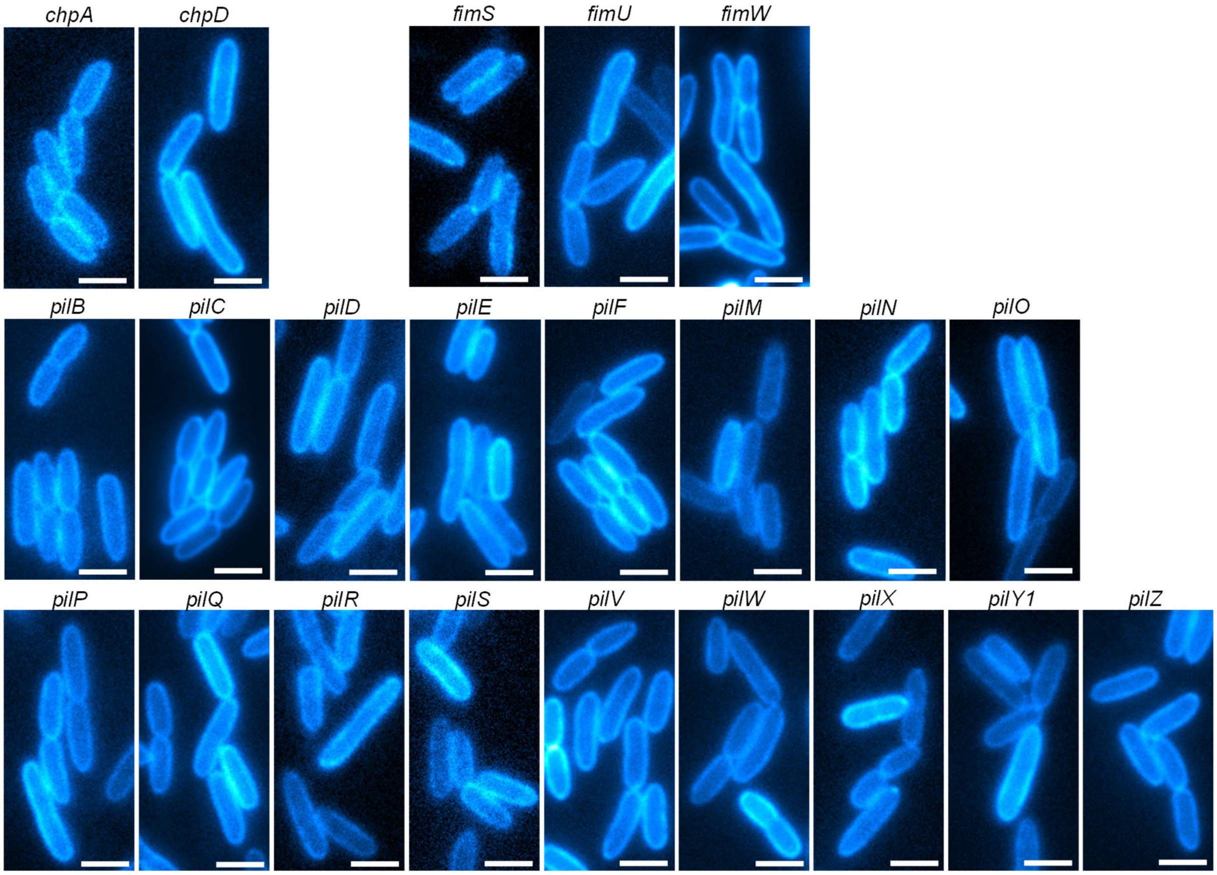

**Supplementary Figure 1. Images of *pilA*-A86C Alexa488-mal labeled cells that did not make any observable pili.** Scale bars = 2 μm.

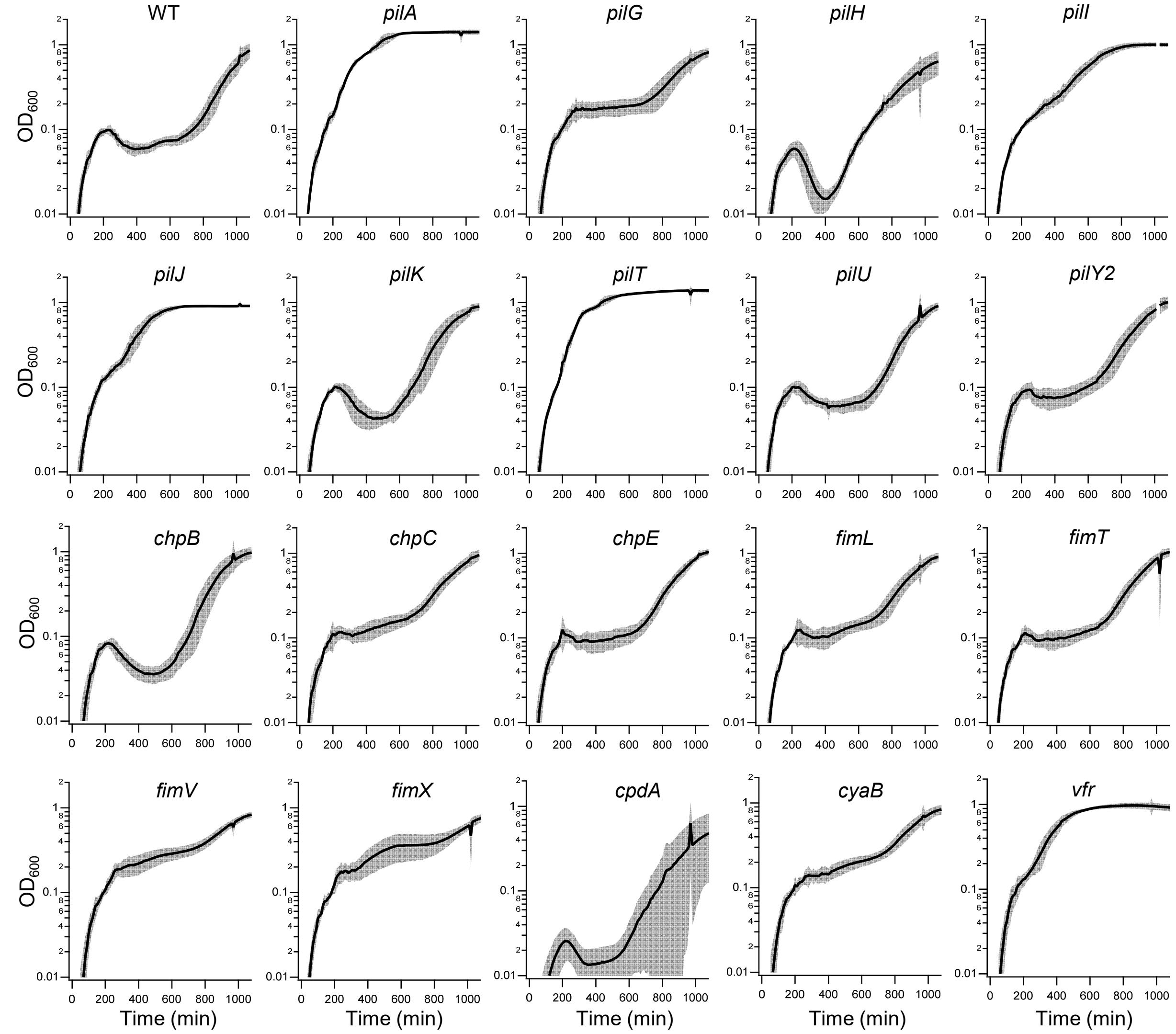

**Supplementary Figure 2. Growth curves of T4P mutants after infection with phage JBD68.** N = 12 experiments (3 biological replicates with 4 technical replicates each). Indicated are the mean (bold line) and standard deviation (shading) at each time point.

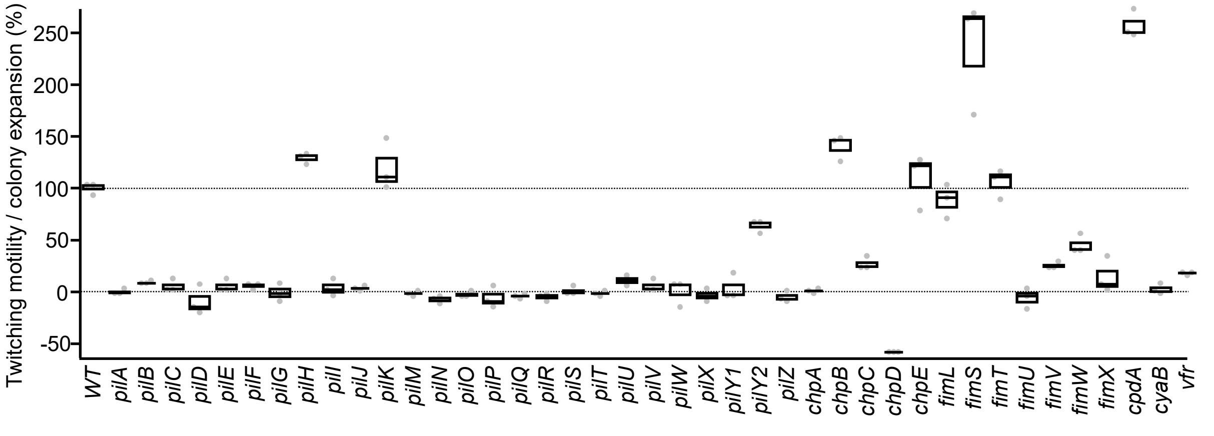

**Supplementary Figure 3. Twitching motility assay for mutants in the extended T4P system.** N = 3 biological replicates were conducted. Twitching motility $T_M$ of mutant M was measured as the size $S_M$ of the stained colony relative to that of WT ($S_{WT}$) and the *pilA* control ($S_{pilA}$) as $T_M = (S_M - S_{pilA}) / (S_{WT} - S_{pilA})$. Negative twitching (e.g. *chpD*) means that the stained colony was smaller than that of *pilA*.

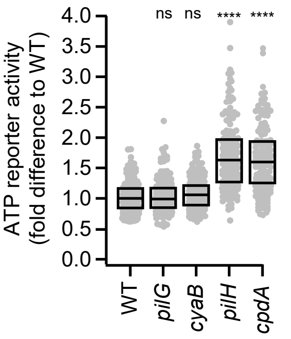

**Supplementary Figure 4. Changes of ATP levels in individual cells in mutants that affect cAMP levels.** N = 150 cells from three biological replicates with 50 cells each were analyzed. **Statistical analysis:** Statistical significance indicates similarity to WT and was tested using a bootstrapped (10,000 iterations) non-parametric Wilcoxon-Mann-Whitney two-sample rank test: (ns) not significant, P > 0.05; (*) P < 0.05; (**) P < 0.01; (***) P < 0.001; (****) P < 0.0001. **Box plots:** Boxes represent median and interquartile range (25th–75th percentiles).

**Strains used in this study**

| MK Strain No. | Genus | Strain | Reference |
|---|---|---|---|
| MK 311 | *E. coli* | WT pEXG2 | [1] |
| MK 315 | *E. coli* | WT pTSN2 | [2] |
| MK 330 | *E. coli* | WT pPaQa | [3] |
| MK 1 | PAO1 | *Pseudomonas aeruginosa*, WT | [4] |
| MK 101 | PAO1 | *pilA*-A86C | [5] |
| MK 43 | PAO1 | *ΔfimU* | [6] |
| MK 44 | PAO1 | *ΔpilE* | [6] |
| MK 45 | PAO1 | *ΔpilV* | [6] |
| MK 46 | PAO1 | *ΔpilW* | [6] |
| MK 47 | PAO1 | *ΔpilX* | [6] |
| MK 30 | PAO1 | *ΔpilB* | [7] |
| MK 31 | PAO1 | *ΔpilH* | [7] |
| MK 29 | PAO1 | *ΔpilA* | [7] |
| MK 36 | PAO1 | *ΔchpA* | [7] |
| MK 37 | PAO1 | *ΔcpdA* | [8] |
| MK 32 | PAO1 | *ΔpilJ* | [3] |
| MK 35 | PAO1 | *ΔpilY1* | [3] |
| MK 38 | PAO1 | *ΔcyaB* | [3] |
| MK 39 | PAO1 | *Δvfr* | [3] |
| MK 163 | PAO1 | *ΔchpB pilA*-A86C | [9] |
| MK 134 | PAO1 | *ΔpilG pilA*-A86C | This Study |
| MK 143 | PAO1 | *ΔpilT pilA*-A86C | This Study |
| MK 144 | PAO1 | *ΔpilU pilA*-A86C | This Study |
| MK 152 | PAO1 | *ΔpilS pilA*-A86C | This Study |
| MK 153 | PAO1 | *ΔpilR pilA*-A86C | This Study |
| MK 162 | PAO1 | *ΔpilK pilA*-A86C | This Study |
| MK434 | PAO1 | *ΔPilN pilA*-A86C | This Study |
| MK436 | PAO1 | *ΔfimV pilA*-A86C | This Study |
| MK495 | PAO1 | *ΔpilF pilA*-A86C | This Study |
| MK497 | PAO1 | *ΔpilD pilA*-A86C | This Study |
| MK501 | PAO1 | *ΔpilO pilA*-A86C | This Study |
| MK503 | PAO1 | *ΔpilM pilA*-A86C | This Study |
| MK505 | PAO1 | *ΔpilP pilA*-A86C | This Study |
| MK507 | PAO1 | *ΔchpD pilA*-A86C | This Study |
| MK509 | PAO1 | *ΔpilQ pilA*-A86C | This Study |
| MK881 | PAO1 | *ΔfimX pilA*-A86C | This Study |

| | | | |
|---|---|---|---|
| MK886 | PAO1 | *ΔfimS pilA*-A86C | This Study |
| MK887 | PAO1 | *ΔfimT pilA*-A86C | This Study |
| MK889 | PAO1 | *ΔfimW pilA*-A86C | This Study |
| MK904 | PAO1 | *ΔpilC pilA*-A86C | This Study |
| MK917 | PAO1 | *ΔpilY2 pilA*-A86C | This Study |
| MK918 | PAO1 | *ΔpilZ pilA*-A86C | This Study |
| MK919 | PAO1 | *ΔchpC pilA*-A86C | This Study |
| MK920 | PAO1 | *ΔpilI pilA*-A86C | This Study |
| MK921 | PAO1 | *ΔfimL pilA*-A86C | This Study |
| MK922 | PAO1 | *ΔchpE pilA*-A86C | This Study |
| MK 124 | PAO1 | *ΔfimU pilA*-A86C | This Study |
| MK 125 | PAO1 | *ΔpilE pilA*-A86C | This Study |
| MK 126 | PAO1 | *ΔpilV pilA*-A86C | This Study |
| MK 127 | PAO1 | *ΔpilW pilA*-A86C | This Study |
| MK 128 | PAO1 | *ΔpilX pilA*-A86C | This Study |
| MK 112 | PAO1 | *ΔpilB pilA*-A86C | This Study |
| MK 113 | PAO1 | *ΔpilH pilA*-A86C | This Study |
| MK 119 | PAO1 | *ΔchpA pilA*-A86C | This Study |
| MK 120 | PAO1 | *ΔcpdA pilA*-A86C | This Study |
| MK 114 | PAO1 | *ΔpilJ pilA*-A86C | This Study |
| MK 115 | PAO1 | *ΔpilY1 pilA*-A86C | This Study |
| MK499 | PAO1 | *ΔcyaB pilA*-A86C | This Study |
| MK 118 | PAO1 | *Δvfr pilA*-A86C | This Study |
| MK752 | PAO1 | *pilB*::*mScarlet*-i3 *pilA*-A86C | [10] |
| MK773 | PAO1 | *pilB*::*mScarlet*-i3 *ΔpilH pilA*-A86C | This Study |
| MK774 | PAO1 | pilB::*mScarlet*-i3 *ΔpilG pilA*-A86C | This Study |
| MK775 | PAO1 | pilB::*mScarlet*-i3 *Δvfr pilA*-A86C | This Study |
| MK 164 | PAO1 | *mRuby3*::*pilT pilA*-A86C | [5] |
| MK776 | PAO1 | *mRuby3*::*pilT Δvfr pilA*-A86C | This Study |
| MK777 | PAO1 | *mRuby3*::*pilT ΔpilG pilA*-A86C | This Study |
| MK778 | PAO1 | *mRuby3*::*pilT ΔpilH pilA*-A86C | This Study |
| MK768 | PAO1 | *pilQ*::*mScarlet*-i3 *pilA*-A86C | This Study |
| MK769 | PAO1 | *pilQ*::*mScarlet*-i3 *ΔpilH pilA*-A86C | This Study |
| MK770 | PAO1 | *pilQ*::*mScarlet*-i3 *ΔpilG pilA*-A86C | This Study |
| MK771 | PAO1 | *pilQ*::*mScarlet*-i3 *Δvfr pilA*-A86C | This Study |
| MK 167 | PAO1 | pPaQa mKate2/YFP | [3] |
| MK 172 | PAO1 | *ΔpilH* pPaQa mKate2/YFP | [3] |
| MK 180 | PAO1 | *Δvfr* pPaQa mKate2/YFP | [9] |
| MK 183 | PAO1 | *ΔcpdA* pPaQa mKate2/YFP | [3] |
| MK 214 | PAO1 | *ΔpilT* pPaQa mKate2/YFP | This Study |

| | | | |
|---|---|---|---|
| MK 215 | PAO1 | *ΔpilU* pPaQa mKate2/YFP | This Study |
| MK 219 | PAO1 | *ΔpilK* pPaQa mKate2/YFP | This Study |
| MK 220 | PAO1 | *ΔchpB* pPaQa mKate2/YFP | [9] |
| MK855 | PAO1 | *ΔchpC* pPaQa mKate2/YFP | This Study |
| MK856 | PAO1 | *ΔchpE* pPaQa mKate2/YFP | This Study |
| MK857 | PAO1 | *ΔcyaB* pPaQa mKate2/YFP | [3] |
| MK858 | PAO1 | *ΔfimL* pPaQa mKate2/YFP | This Study |
| MK859 | PAO1 | *ΔfimV* pPaQa mKate2/YFP | This Study |
| MK860 | PAO1 | *ΔpilI* pPaQa mKate2/YFP | This Study |
| MK861 | PAO1 | *ΔpilJ* pPaQa mKate2/YFP | [3] |
| MK862 | PAO1 | *ΔpilY2* pPaQa mKate2/YFP | This Study |
| MK905 | PAO1 | *ΔfimX* pPaQa mKate2/YFP | This Study |
| MK906 | PAO1 | *ΔfimT* pPaQa mKate2/YFP | This Study |
| MK907 | PAO1 | *ΔpilG* pPaQa mKate2/YFP | This Study |
| MK 575 | PAO1 | *pilA*-A86C pAY9 (ATP biosensor) | [10] |
| MK 913 | PAO1 | *ΔcpdA pilA*-A86C pAY9 (ATP biosensor) | This Study |
| MK 914 | PAO1 | *ΔcyaB pilA*-A86C pAY9 (ATP biosensor) | This Study |
| MK 915 | PAO1 | *ΔpilH pilA*-A86C pAY9 (ATP biosensor) | This Study |
| MK 916 | PAO1 | *ΔpilG pilA*-A86C pAY9 (ATP biosensor) | This Study |

**Primers used in this study**

| **Primer** | **Sequence (5' to 3')** | **Referen ce** |
|---|---|---|
| pEXG2_Ver1 | GTTGCATGGGCATAAAGTTGCC | [5] |
| pEXG2_Ver2 | CGGGTCCTCAACGACAGG | [5] |
| pTN7_Ver3 | TTATCTGGTTGGCCTGCAAGG | [5] |
| pTN7_Ver4 | CGATCCATTGCTGTTGACAAAGG | [5] |
| pilG_P1 | GATACAAAGCTTCGTTGAAGACCTTCAGCGGG | |
| pilG_P2 | CGGATATCAGGAAACGGCGTCGGATTGCTGTTCCA TGTTCGC | |
| pilG_P3 | GCGAACATGGAACAGCAATCCGACGCCGTTTCCT GATATCCG | |
| pilG_P4 | GATACAAAGCTTGGTCAGGGTGAGCGATGC | |
| pilT_P1 | GGAAGCATAAATGTAAAGCACCAGTTCGGCCTGCT TGC | |
| pilT_P2 | CCAGGATCAGAAGTTTTCCGGCTCGGTAATATCCAT GGGACTCC | |
| pilT_P3 | GGAGTCCCATGGATATTACCGAGCCGGAAAACTTC TGATCCTGG | |
| pilT_P4 | GAGTCGACCTGCAGATTCAGCTCTTCCAGGGTCG | |

| | |
|---|---|
| pilU_P1 | GGAAGCATAAATGTAAAGCACGATGCTCTCCGAGTCGC |
| pilU_P2 | GGTTCAGCGGAAGCGCCGCTTTTCGAATTCCATGATGTTCTCGC |
| pilU_P3 | GCGAGAACATCATGGAATTCGAAAAGCCGGCGCTTCCGCTGAACC |
| pilU_P4 | GAGTCGACCTGCAGAGCCTACCTGGAGGTCTGG |
| pilS_P1 | GATACAAAGCTTTGACCTTCGGTCTCTTCGAC |
| pilS_P2 | CTTCCGTCAGCTGAGTTTGCGAGCCGTTCAGCGCGCACGG |
| pilS_P3 | CCGTGCGCGCTGAACGGCTCGCAAACTCAGCTGACGGAAG |
| pilS_P4 | GATACAAAGCTTCCGATCTGGTTGCGCAGG |
| pilR_P1 | GATACAAAGCTTGACGAATACCCCGGCAGG |
| pilR_P2 | CACTTTCAGTCGATGCCCAGTTTTTGTCGGCTCATGCGTGC |
| pilR_P3 | GCACGCATGAGCCGACAAAAACTGGGCATCGACTGAAAGTG |
| pilR_P4 | GATACAAAGCTTCCGATCTGGTTGCGCAGG |
| pilK_P1 | GGAAGCATAAATGTAAAGCAGCCGCGGCTTCGC |
| pilK_P2 | ACCGGCTCGCCGTTCGCCTGCATGC |
| pilK_P3 | GAACGGCGAGCCGGTCGCC |
| pilK_P4 | GAGTCGACCTGCAGAGAGTTCGTCGGTGCGCAG |
| pilN_P1 | GATACAAAGCTTTTGACCGTGGCGGTAGTG |
| pilN_P2 | GACTCATTTCTTGGCTCCGTTGATCCGTGCCATCAG |
| pilN_P3 | CTGATGGCACGGATCAACGGAGCCAAGAAATGAGTC |
| pilN_P4 | GATACAAAGCTGGACACGCCGCTGACGAAG |
| fimV_P1 | GATACAAAGCTTAACGTTACCTGCATCCAGG |
| fimV_P2 | GGATCAGGCCAGGCGCTCACGAAGCCGAACCATAGTG |
| fimV_P3 | CACTATGGTTCGGCTTCGTGAGCGCCTGGCCTGATCC |
| fimV_P4 | GATACAAAGCTTATCCGCCCGATATCCAGC |
| pilF_P1 | GATACAAAGCTTGCACCGAACGACGAGTTG |
| pilF_P2 | TTCATCATTTTTCCGCCTGGGCGCGTACAGTCATCGG |
| pilF_P3 | CCGATGACTGTACGCGCCCAGGCGGAAAAATGATGAA |
| pilF_P4 | GATACAAAGCTTTATGGATCTCGGTGGTGCC |
| pilD_P1 | GATACAAAGCTTACGCTCGAAAAAATTCCGC |
| pilD_P2 | GGGTCATTTGAATCCGGCGTCGAGGAGGGGCATCAG |

| | |
|---|---|
| pilD_P3 | CTGATGCCCCTCCTCGACGCCGGATTCAAATGACCC |
| pilD_P4 | GATACAAAGCTTCTGGGCCTGGAGGATCGA |
| pilO_P1 | GATACAAAGCTTTGCGCTGGTATTCCTTGG |
| pilO_P2 | CTCTCATTTCTTCAGCCCACTGGCCAGACTCATTTC |
| pilO_P3 | GAAATGAGTCTGGCCAGTGGGCTGAAGAAATGAGAG |
| pilO_P4 | GATACAAAGCTTGACGCTCCAGCCAGTTCC |
| pilM_P1 | GATACAAAGCTTATTAGGCTTTTCACATCGAC |
| pilM_P2 | CCATCAGTCGAAACTCCTTATGAGCCCTAGCACGACC |
| pilM_P3 | GGTCGTGCTAGGGCTCATAAGGAGTTTCGACTGATGG |
| pilM_P4 | GATACAAAGCTTCTGGGTCACCGCCTTGAC |
| pilP_P1 | GATACAAAGCTTTGGGCTACAACTTCCATCTG |
| pilP_P2 | CCCTCAGGAGCGTTCCTTCAGGCGGGCTCTCATTTC |
| pilP_P3 | GAAATGAGAGCCCGCCTGAAGGAACGCTCCTGAGGG |
| pilP_P4 | GATACAAAGCTTGGGCTTGCCTTGACCGGA |
| chpD_P1 | GATACAAAGCTTTGGCATGAGCCGGAACAC |
| chpD_P2 | AGTTCAGCCGCGCACGGCCTGCAGGCCGGCCATGGA |
| chpD_P3 | TCCATGGCCGGCCTGCAGGCCGTGCGCGGCTGAACT |
| chpD_P4 | GATACAAAGCTTGTCGACGATCCCCGCCAG |
| pilQ_P1 | GATACAAAGCTTTACATGGACGAGGTTCGCG |
| pilQ_P2 | GATATCAGCGACCGATTGCGAGGCCACTGTTCATCGTC |
| pilQ_P3 | GACGATGAACAGTGGCCTCGCAATCGGTCGCTGATATC |
| pilQ_P4 | GATACAAAGCTTAATTTCCTGGACGACCAGG |
| fimX_P1 | GATACAAAGCTTTCCGCCAGGTACGCAATG |
| fimX_P2 | TCTTCATTCGTCTCCCGACTTTTCGATGGCCATGGA |
| fimX_P3 | TCCATGGCCATCGAAAAGTCGGGAGACGAATGAAGA |
| fimX_P4 | GATACAAAGCTTTTCGCCGAACTCAAGCGC |
| fimS_P1 | GGAAGCATAAATGTAAAGCACTGCCGGCTTCGATTTC |
| fimS_P2 | GCTTCCTGCATGAATCGGATAGG |
| fimS_P3 | CTATCCGATTCATGCAGGAAGCC |
| fimS_P4 | GAGTCGACCTGCAGACAGCTCGATCCCCTTGCG |
| fimT_P1 | GGAAGCATAAATGTAAAGCACGTGCCCACTGCAGTGCC |

| | |
|---|---|
| fimT_P2 | GAAGTGCTCGACCTTTCGACCA |
| fimT_P3 | GAAAGGTCGAGCACTTCCGGAT |
| fimT_P4 | GAGTCGACCTGCAGAAGGTCAGATGTTCCACGG |
| fimW_P1 | GGAAGCATAAATGTAAAGCAAAGGCGCCGAGC |
| fimW_P2 | GACTTCCAGCTCTGGTTTTCCATG |
| fimW_P3 | GGAAAACCAGAGCTGGAAGTCTCTGTAGA |
| fimW_P4 | GAGTCGACCTGCAGAGCGCGCGCGGCGTTCG |
| pilC_P1 | GGAAGCATAAATGTAAAGCAATCATCGCCCAGCG |
| pilC_P2 | GACGACGTTGGCCTTCACCA |
| pilC_P3 | GAAGGCCAACGTCGTCGGAT |
| pilC_P4 | GAGTCGACCTGCAGAGGTGATCGGCATCG |
| pilY2_P1 | GATACAAAGCTTTCGCACCATTCGCATCGC |
| pilY2_P2 | TCATCGGGGCTGCTCCTCAGGCAGCACTTTCATAT CAGT |
| pilY2_P3 | ACTGATATGAAAGTGCTGCCTGAGGAGCAGCCCC GATGA |
| pilY2_P4 | GATACAAAGCTTAAGGAACTGCGGGTCAAGG |
| pilZ_P1 | GATACAAAGCTTGGCCCTGGAGGAACTGTTG |
| pilZ_P2 | GCGTTACATCGTGTGGGTGGGTGGCAAACTCATGC |
| pilZ_P3 | GCATGAGTTTGCCACCCACCCACACGATGTAACGC |
| pilZ_P4 | GATACAAAGCTTACTGGCTTGGCGGTGATC |
| chpC_P1 | GATACAAAGCTTTCTATGCCCAGCACATCGATG |
| chpC_P2 | GGCTCAGATCAGGCCGGCCACGGCCTGGTTCATG TTT |
| chpC_P3 | AAACATGAACCAGGCCGTGGCCGGCCTGATCTGA GCC |
| chpC_P4 | GATACAAAGCTTCAGGATCGGCCCTCTGCG |
| pilI_P1 | GATACAAAGCTTACGTACCCAGCTTCACCC |
| pilI_P2 | GCTGTTTATACGGCGACGTCCTGAACGTCCGACAT GCG |
| pilI_P3 | CGCATGTCGGACGTTCAGGACGTCGCCGTATAAAC AGC |
| pilI_P4 | GATACAAAGCTTGTCTGTTCGCTGGCGAGG |
| fimL_P1 | GATACAAAGCTTGCCATCGACCTGGTCTCG |
| fimL_P2 | CCATCAGGCGGCCACCGGGGCTCCTGTGACCATC GG |
| fimL_P3 | CCGATGGTCACAGGAGCCCCGGTGGCCGCCTGAT GG |
| fimL_P4 | GATACAAAGCTTCGCCAGCAACACCTCGTC |
| chpE_P1 | GGAAGCATAAATGTAAAGCACCAGCGACCGCTTCG |
| chpE_P2 | CCGCGCAGGAAGATGGCGAGCATGGAACG |
| chpE_P3 | CATCTTCCTGCGCGGGCTGTAGAC |
| chpE_P4 | GAGTCGACCTGCAGACTGCCTGCTCTCCCGC |
| FTL1_FP.For | GCCGGCGGGTCGGGC |

| | |
|---|---|
| FTL1_FP.Rev | ACCACCGGCACCACCCG |
| pilQ_FTL1_CT_F1.For | GGAAGCATAAATGTAAAGCACTGCAACTGTCGGCGATGG |
| pilQ_FTL1_CT_F1.Rev | CCCGACCCGCCGGCGCGACCGATTGCGATGGC |
| pilQ_FTL1_CT_F2.For | GGTGGTGCCGGTGGTTGATATCGAACAACGTGGTCAACC |
| pilQ_FTL1_CT_F2.Rev | GAGTCGACCTGCAGAGGCCTTTCGTCGGTCTCC |

### Supplementary References

1. Hmelo, L.R., et al., *Precision-engineering the Pseudomonas aeruginosa genome with two-step allelic exchange.* Nat Protoc, 2015. **10**(11): p. 1820-41.
2. Choi, K.H., et al., *A Tn7-based broad-range bacterial cloning and expression system.* Nat Methods, 2005. **2**(6): p. 443-8.
3. Persat, A., et al., *Type IV pili mechanochemically regulate virulence factors in <i>Pseudomonas aeruginosa</i>.* Proceedings of the National Academy of Sciences, 2015. **112**(24): p. 7563-7568.
4. Jacobs, M.A., et al., *Comprehensive transposon mutant library of Pseudomonas aeruginosa.* Proc Natl Acad Sci U S A, 2003. **100**(24): p. 14339-44.
5. Koch, M.D., et al., *Competitive binding of independent extension and retraction motors explains the quantitative dynamics of type IV pili.* Proceedings of the National Academy of Sciences, 2021. **118**(8): p. e2014926118.
6. Marko, V.A., et al., *Pseudomonas aeruginosa type IV minor pilins and PilY1 regulate virulence by modulating FimS-AlgR activity.* PLoS Pathog, 2018. **14**(5): p. e1007074.
7. Bertrand, J.J., J.T. West, and J.N. Engel, *Genetic analysis of the regulation of type IV pilus function by the Chp chemosensory system of Pseudomonas aeruginosa.* J Bacteriol, 2010. **192**(4): p. 994-1010.
8. Inclan, Y.F., M.J. Huseby, and J.N. Engel, *FimL Regulates cAMP Synthesis in Pseudomonas aeruginosa.* PLOS ONE, 2011. **6**(1): p. e15867.
9. Koch, M.D., et al., *<i>Pseudomonas aeruginosa</i> distinguishes surfaces by stiffness using retraction of type IV pili.* Proceedings of the National Academy of Sciences, 2022. **119**(20): p. e2119434119.
10. Yusuf, A.O., Z.K. Modi, and M.D. Koch, *Rapid activation of dormant type IV pili enables a dispersal–infection tradeoff in environments with fluctuating nutrients.* bioRxiv, 2026: p. 2026.04.23.720421.
